# Collective Posterior Inference from Highly Variable Empirical Replicates

**DOI:** 10.64898/2026.01.26.701673

**Authors:** Nadav Ben Nun, Saharon Rosset, David Gresham, Yoav Ram

**Affiliations:** School of Zoology, Faculty of Life Sciences, Tel Aviv University, Tel Aviv, Israel; Edmond J. Safra Center for Bioinformatics, Tel Aviv University, Tel Aviv, Israel; Department of Statistics, Tel Aviv University, Tel Aviv, Israel; Department of Biology, Center for Genomics and Systems Biology, New York University, New York, United States

## Abstract

High-throughput experimental platforms now routinely generate data from dozens or hundreds of independent observations. Simulation-based inference (SBI) offers a powerful framework for estimating model parameters from such complex datasets, but standard methods struggle to scale to the noisy multiple-replicates regime without incurring prohibitive computational costs or careful hyperparameter tuning.

Here, we introduce a new method for fast and robust collective posterior inference from multiple independent replicates using a robust product-of-experts aggregation scheme that automatically mitigates the influence of outliers. Evaluating it on synthetic and empirical evolutionary datasets, we find it achieves state-of-the-art estimation accuracy and computational efficiency, including inference from noisy observations. Our method is compatible with any SBI framework, providing a scalable, plug-and-play solution for inference from noisy multiple-replicate datasets.

## 1. Introduction

### Bayesian inference from multiple replicates

Many biological studies aim to infer parameters of a generative model from data while quantifying uncertainty. In a Bayesian framework, this uncertainty is captured by a posterior distribution over model parameters conditioned on the data. Increasingly, experiments produce multiple replicates collected under the same protocol and intended to reflect a shared underlying process. Here, a *replicate* is one observation (possibly multivariate, e.g., time series of genotype frequencies); *multiple replicates* are a set of replicates assumed to be conditionally independent given the model parameters, that is, independent draws from the same generative model with shared parameter values.

In practice, however, departures from this assumption are common: technical replicates can differ due to variability in experimental procedures and batch effects (1–3), biological replicates capture heterogeneity across individuals or populations (4,5), and faulty or mislabeled samples can introduce additional inconsistencies. The central problem we address here is how to construct a posterior distribution that reflects the shared signal from multiple replicates while remaining robust to departures from the assumed generative model.

### Simulation-based inference

Classical Bayesian inference requires an explicit likelihood function that specifies the probability of the data given the model parameters. For many scientific models, however, this likelihood is intractable (6–10). To address this*, simulation-based inference* (SBI; also *likelihood-free inference*) is often used. Initially developed to study human evolution (11,12), SBI has been adopted across various scientific domains, including experimental evolution (13–15), economics (16,17), epidemiology (18,19), and astronomy (20–22).

The simplest SBI method is *Rejection Approximate Bayesian Computation* (rejection-ABC) (23). Given a set of empirical observations, the analyst defines (i) a (stochastic) generative model, implemented via a simulator, that maps parameters to a synthetic observation; (ii) a prior distribution over the model parameters; (iii) a summary statistics of observations; (iv) a function that measures similarity between two summary-statistics; and (v) a similarity threshold. Parameter values drawn from the prior are accepted as posterior samples if, when used to run the simulator, they generate a synthetic observation whose summary statistics are within the similarity threshold of the empirical summary statistics. More efficient ABC variants, such as MCMC-ABC and SMC-ABC, improve sample efficiency by adaptively exploring the parameter space and reducing the number of required simulations.

More recently, neural density estimators – artificial neural networks that approximate complex probability distributions – have been integrated with SBI (24). For example, in *Neural Posterior Estimation* (NPE) (25,26), a training set is generated by sampling parameters from the prior distribution and using a simulator to produce corresponding synthetic observations. A neural density estimator—an artificial neural network that outputs the parameters of a flexible conditional density—is then trained on parameter-observation pairs to learn an approximation to the posterior distribution. Once trained, the estimator can be conditioned on real observations to approximate the posterior distribution, enabling posterior evaluation at specific parameter values and efficient posterior sampling. However, these methods are typically designed to condition on a single replicate, whereas many experimental designs collect multiple replicates to capture biological variability and to overcome technical measurement error.

### SBI with multiple replicates

Although SBI is widely used, comparatively few methods directly target posterior inference from multiple replicates. With an explicit likelihood, the posterior conditioned on all replicates is obtained by factoring the likelihood across replicates and combining it with the prior. In likelihood-free settings, this route is unavailable, and conditioning on multiple replicates is less straightforward.

Several recent approaches extend SBI to inference from multiple replicates. We categorize these methods by their mechanism for aggregating information and their specific limitations in terms of computational efficiency and model flexibility.

### Embedding-based approaches (NPE+PIE)

This approach is inspired by exchangeable neural-network architectures (28,29) and implemented in the *sbi* framework (27). A permutation-invariant embedding (PIE) network learns embeddings for single replicates and then performs a permutation-invariant aggregation (e.g., mean or sum) on these embeddings. The aggregated embedding serves as input to the density estimator. in a recent practical guide to SBI (30), NPE+PIE was highlighted as an approach for neural posterior estimation from multiple replicates.

Training of NPE+PIE requires a large training set, because the PIE network and the density estimator are trained together on randomly constructed subsets of replicates. Furthermore, it enforces a rigid simulation structure: to learn the embeddings, the training data must capture the variance of the simulator for fixed parameters, often requiring batches of replicates for every θ rather than independent (x, θ) pairs.

### Hierarchical inference approaches

Hierarchical NPE (HNPE) explicitly models the hierarchical data-generating process (31). It learns separate estimators for replicate-specific and shared parameters, allowing it to disentangle sources of variation. However, like NPE+PIE, HNPE requires specialized, hierarchical neural architectures and relies on the simulator ability to generate data with a specific hierarchical structure, which limits its applicability to generic simulators and noise structures. It also imposes a significant simulation burden, as the network learns not just the shared parameters but also the distribution of replicate-specific latent parameters, requiring a large dataset for every training example to characterize the hierarchical structure.

Another hierarchical approach assumes that each replicate has its own latent parameters, which are sampled from a shared distribution (32), as in standard random-effects models. Replicate-specific posteriors are amortized with conditional normalizing flows, and inference over the parameters of the shared distribution combines a mixed-effect model with Monte Carlo sampling from these learned individual posteriors. The second step can be slow, especially when replicates are noisy, and the replicate-specific posteriors are broad, requiring many samples for stable estimates.

### Score-based and diffusion approaches

Recent work has adapted score-based generative models to the multiple-replicate setting (33–35). These approaches typically learn the gradient of the log-posterior (the score) or a flow-matching objective conditioned on single replicates. While highly accurate, they introduce significant inference complexity. Methods like NPSE (33,34) rely on iterative Langevin dynamics, requiring hundreds of network evaluations per sample. Similarly, the recent flow-matching approach of Arruda et al. (35) requires careful differential equation solver scheduling and error damping to avoid numerical instabilities during the integration of the flow.

### Our contribution

Here, we propose a novel method for simple and flexible inference from multiple replicates, the *collective posterior*, that combines posteriors conditioned on individual replicates into a single posterior conditioned on the full dataset. Unlike existing SBI pipelines, our method allows that some replicates may not have been generated by the same generative model as the others; the method can downweight or even effectively ignore replicates whose individual posteriors are inconsistent with the dominant generative model. This feature is crucial in applied settings: empirical datasets routinely include faulty measurements, mislabeled samples, or biological outliers that pass standard quality-control filters but distort inference if treated on equal footing with the rest of the data. Our method yields posterior estimates that remain stable in the presence of such aberrant observations, providing a principled safeguard against the bias these observations can introduce. The collective posterior is general and integrates with existing Bayesian inference frameworks, from rejection-ABC to neural SBI, and also with likelihood-based inference. We demonstrate the collective posterior on several simulation-based inference tasks, using both rejection-ABC and NPE as individual posterior estimators, and compare it to a prominent existing approach, NPE+PIE. We also applied it to quantify how genetic background affects the evolutionary dynamics of *GAP1* copy-number variants (CNVs) (36) and CNV stability (37) in *Saccharomyces cerevisiae*.

## 2. Methods

### Existing Methods

#### Rejection-ABC

Rejection-Approximate Bayesian Computation (23) is a simple SBI method. It includes the following steps: i) Sample parameter proposals from the prior distribution, θ_i_∼p(θ). ii) Simulate synthetic data, x_i_∼f(θ_i_), where f is a simulator that generates data from the intractable likelihood, p(x|θ). iii) Compute summary statistics for the synthetic data, s(x_i_), and empirical data, s(x_o_). iv) Accept proposal θ_i_ if d(s(x_o_),s(x_i_)) ≤ ε, where d(⋅,⋅) is a dissimilarity function and ε is a dissimilarity threshold. v) Repeat until enough proposals are accepted. With sufficient summary statistics and ε→0, the distribution of accepted parameters converges to p(θ|x_o_). This approach can be easily parallelized but can suffer from low acceptance rates (and therefore low computational efficiency) when the prior and posterior distributions have a small overlap, when the number of parameters is large, and when ε is small.

#### Neural Posterior Estimation (NPE)

A prominent example of SBI with neural density estimation is *Neural Posterior Estimation* (25,26). NPE directly approximates the posterior density of model parameters conditioned on an observation, p(θ|x) (Figure S1a). The neural conditional density estimator q_φ_(θ|x) (parameterized by φ) is trained to approximate p(θ|x): a training set of pairs (θ_i_, x_i_) is generated by sampling parameters from the prior, θ_i_∼p(θ), and simulating observations, x_i_∼f(θ_i_); then gradient-based optimization is used to find φ* that maximizes Σ_i_log(q_φ_(θ_i_|x_i_)). This yields an *amortized* posterior estimator, q_φ*_(θ|x), which, when conditioned on a specific observation x_o_, provides q_φ*_(θ|x_o_)≈p(θ|x_o_). Importantly, once trained, evaluating the posterior estimator q_φ*_ on a new observation x_o_ does not require additional training or simulation, in contrast to sampling methods such as rejection-ABC, which require additional simulations to infer the posterior distribution for every new observation. The neural density estimator can be implemented by a normalizing flow, which provides a general way of constructing flexible probability distributions over continuous random variables (38). Our previous studies (39,40) demonstrated that NPE infers more accurate and confident posterior distributions from evolutionary experimental data than popular sampling-based Bayesian inference methods (rejection-ABC and SMC-ABC (41)).

#### Neural Posterior Score Estimation (NPSE)

Recent studies on SBI from independently and identically distributed (i.i.d.) observations have introduced score-based models for improved accuracy of SBI (33–35). While their results are promising, we did not observe any advantage in using NPSE over NPE for our empirical data (Figure S6). We therefore proceeded with NPE as the neural SBI method for individual posterior estimation. For tasks where NPSE is better at individual posterior estimation, our method can be applied using NPSE as the individual posterior estimator, or by leveraging NPSE’s i.i.d. formulation (33,34).

### Collective posterior distribution

#### Standard collective posterior distribution

Assuming that multiple replicates {x_i_ }^n^_i=1_ are independent conditioned on the model parameters, θ, that is, p(x_i_,x_j_|θ) = p(x_i_|θ)p(x_j_|θ), we can compute the *collective* posterior distribution of model parameters conditioned on all replicates, p(θ|x_1_,…,x_n_), from individual posterior distributions, p(θ|x_i_), and the prior distribution, p(θ), by applying Bayes’ theorem twice (a full derivation is available in the SI),

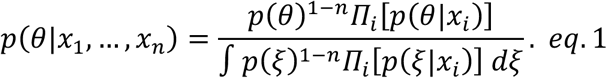

#### Robust posterior distribution

Posterior distribution approximation in SBI, while accurate in the high posterior density region (HDR), tends to be uncalibrated in low-density regions, sometimes producing erratic posterior density values (Table S3). To mitigate this, we introduce a robust individual posterior distribution,

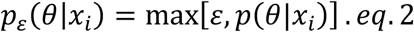

We combine these robust individual posterior distributions into a robust collective posterior distribution,

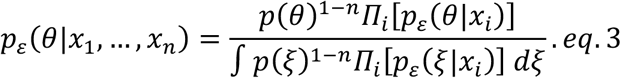

For ε→0, we have p_ε_(θ|x_i_)→p(θ|x_i_) and and p_ε_(θ| x_1_,…,x_n_)→p(θ|x_1_,…,x_n_). High values of ε will focus only on high-density regions of the individual posterior distributions, overfitting to a given replicate set. An optimal choice of ε balances robustness to outliers with accurate parameter inference.

#### Sampling from the robust collective posterior distribution

To sample from the collective posterior (Eq. 3), we implemented and compared three distinct sampling strategies. Based on the trade-off between computational efficiency and approximation accuracy, *Sampling Importance Resampling (SIR)* was selected as the primary method for the results presented in this study. The other strategies are detailed in the supplementary information.

#### Sampling Importance Resampling (SIR)

To efficiently approximate the collective posterior without relying on sequential chains (i.e., MCMC), we used an SIR scheme (42). We used the prior distribution p(θ) as the proposal distribution to ensure robust coverage of the parameter space while not relying on the sampling efficiency of the individual posterior distribution. Importance weights were calculated as the ratio of the unnormalized robust collective posterior density to the prior density. To counteract the artificial variance reduction inherent in the product-of-experts aggregation, where multiplying independent densities shrinks the standard deviation by a factor of approximately 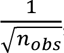, we applied temperature scaling to the importance weights. Then, at resampling, we applied Gaussian jittering to the results. We set the temperature 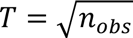 to prevent weight collapse. This vectorized approach enabled rapid, high-throughput posterior approximation.

### Test cases

#### Synthetic evolutionary data

Consider a Wright–Fisher model of a haploid population of fixed size N=10^4^ with three bi-allelic loci. The eight genotypes are represented by a bit-string (b_1_,b_2_,b_3_)∈{0,1}^3^, with the population initialized as all wild type, (0,0,0). Genotype fitness is multiplicative: the presence of a mutant allele 1 contributes a fitness factor 1+s_i_, so the fitness of genotype g = (b_1_,b_2_,b_3_) is ∏^3^_i=1_[(1+s_i_)^bi^].

Mutation is independent across loci. By default, we allow only forward mutation 0→1 with per-locus rates μ_1_, μ_2_, μ_3_, so that mutation is applied by an 8×8 transition matrix M, where M[g,h] is the probability of mutation from genotype g to genotype h; this probability factorizes across loci by the mutation rates. Genetic drift arises from multinomial sampling given the expected genotype frequencies after mutation and selection.

We simulate the model for 1000 generations, and we record allele (rather than genotype) frequencies every 100 generations. The simulator returns the time points and three allele-frequency time series as a single 30-dimensional vector, which is used as a single replicate.

For testing, we generate a dataset of 200 observations, each consisting of 10 replicates (Figure 2). The intrinsic stochoasticity of this model produces substantial variance among replicates. However, to better resemble real datasets, we further contaminate the data: of the 10 replicates, seven replicates correspond to simulations using parameters perturbed by additive Gaussian noise sampled from N(0, 0.1), mimicking biological variability through a hierarchical process. The remaining three are simulations of the same parameters, perturbed by a large additive Gaussian noise sampled from N(0,1), which introduces outlier observations to the dataset. Finally, to simulate measurement error, we perturb each observation with heteroscedastic Gaussian noise, where the standard deviation is set to 10% of the observed value (σ = 0.1⋅x_o_). Overall, this design aims to mimic empirical datasets, which are inherently noisy due to both stochastic and measurement variability. We found that introducing these perturbations produces datasets that closely resemble experimental observations in their variability and structure (Figure S4).

**Figure 1.**
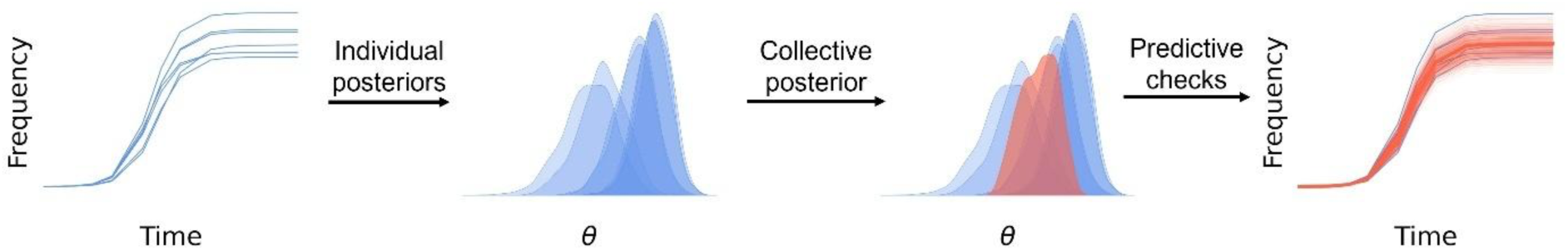
Collective posterior inference. From left to right: independent replicates (e.g., time series; blue) are analyzed to obtain replicate-specific posterior distributions (blue), which are then combined to form a collective posterior conditioned on the full replicate set (red). Finally, posterior predictive checks assess whether simulations from the collective posterior (red) reproduce the empirical distribution of time series across replicates (blue), including between-replicate variability.

**Figure 2.**
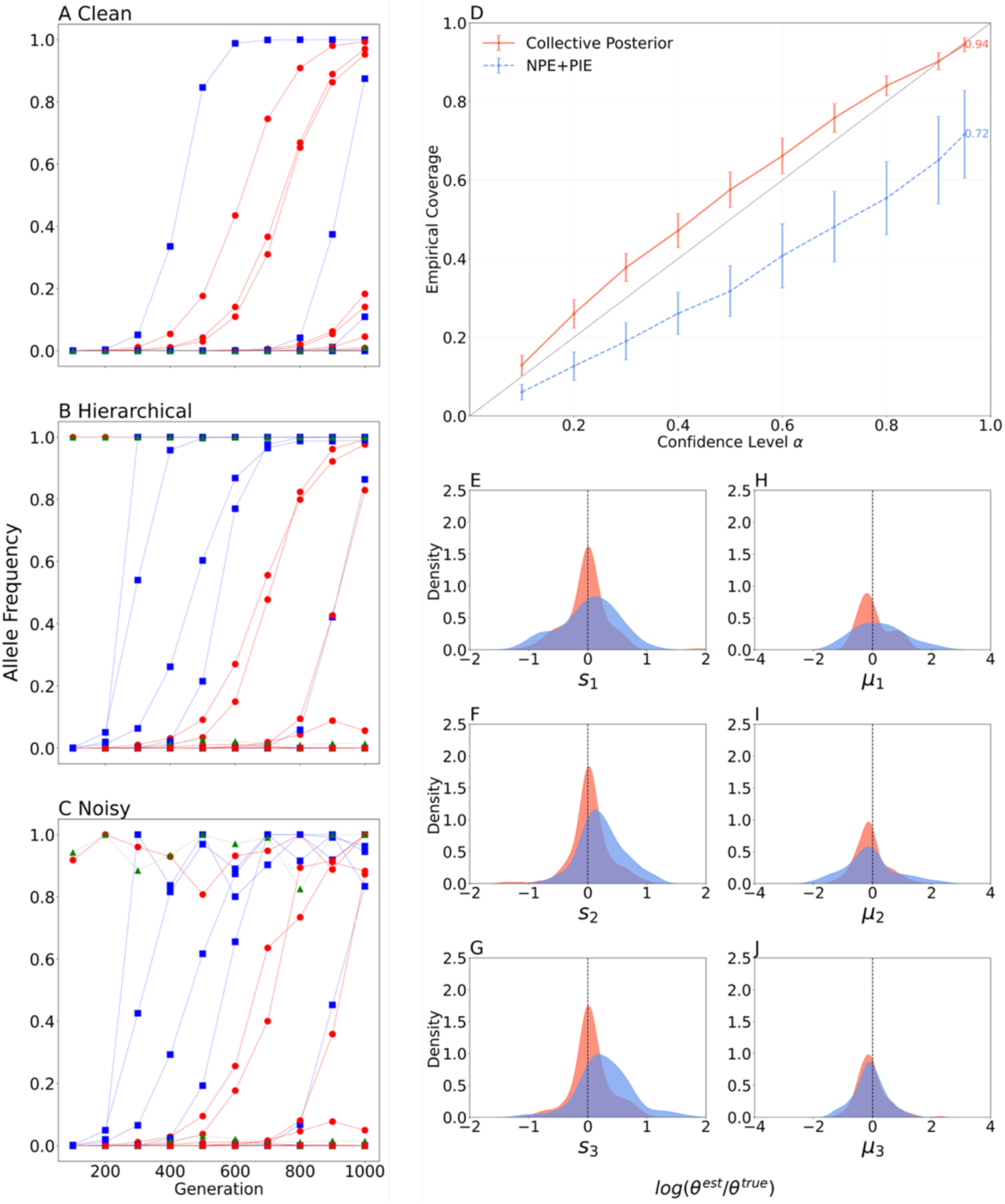
The collective posterior outperforms NPE+PIE on noisy evolutionary simulations. (A-C) Synthetic replicate sets. Each test instance comprises n=10 Wrigh-Fisher replicate time series simulated with shared parameters, θ (A, Clean). To introduce between-replicate heterogeneity, we perturb θ using for 7 replicates with moderate Gaussian noise and for 3 replicates with larger perturbations (B, Hierarchical). Finally, we add measurement noise to the time series (C, Noisy). **(D) Coverage.** For each nominal confidence level α, we compute the empirical coverage Pr[θ_i_∈HDR_α_] of per-parameter highest-density regions (ideal: diagonal). The collective posterior achieved consistently better coverage (closer to the diagonal) than NPE+PIE, indicating less overconfidence. **(E-J) Accuracy.** Distributions of per-parameter estimation errors (defined as the log-ratio of the posterior mean and true value). Across parameters, the collective posterior is more tightly centered around zero, indicating more accurate inference.

#### Empirical evolutionary datasets

We consider our recent study (36) in which we inferred the parameters of a Wright-Fisher model from experimental evolutionary data. The data are time series of the proportions of cells with more than one copy of the *GAP1* gene in haploid yeast populations (4 different genotypes, 27 time series in total) in a glutamine-limited environment over 116 generations. By inserting a green fluorescent protein (GFP) next to *GAP1*, the CNV proportion was monitored throughout the experiment using flow cytometry (43). The inference task involves estimating three parameters: the selection coefficient s of *GAP1* duplications, its formation rate δ, and its initial undetectable frequency φ, which is non-zero due to cells with non-GFP *GAP1* duplications in the founder population. The evolutionary model includes four genotypes: WT, with an initial proportion X^0^_WT_ = 1-φ and fitness of w_WT_=1; C^-^, representing cells with *GAP1* duplications that formed before the fluorescent reporter insertion (thus undetectable throughout the experiment), with an initial proportion X^0^_C-_= φ and fitness of w_C-_ = 1+s; C^+^, representing cells with *GAP1* duplications that formed after reporter insertion, with an initial proportion X^0^_C+_ = 0 and fitness w_C+_ = 1+s; and B, representing other beneficial genotypes, with initial proportion X^0^_B_ = 0 and fitness w_B_ = 1.001. At generation *t*, mutations occur with rates δ from *WT* to *C^+^* and μ=10^-5^ from *WT* to *B*, so that the change in the genotype frequencies is described by

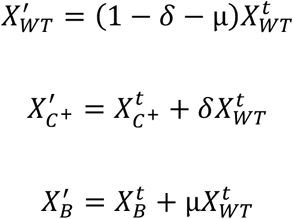

Then, genotype frequencies X_i_’ change following their associated fitness w_i_,

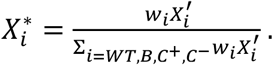

Finally, frequencies change due to random genetic drift,

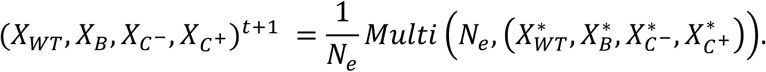

The effective population size in the chemostats was estimated to be N_e_=3.3×10^8^ (Avecilla et al. 2022). Here, we use a lower value of N_e_=10^7^ for training and testing the neural density estimator to increase the variance between observations, posing a more difficult inference challenge. Our main analyses focus on the empirical evolutionary data, and we also benchmarked performance on synthetic datasets by simulating 500 observations, each consisting of 10 replicates (see SI for results).

#### Swamp sparrow song dataset

Lachlan et al. (5) analyzed song patterns from six geographically distinct swamp sparrows (*Melospiza georgiana*) populations to infer the parameters of a cultural evolutionary model of syllables transmission: μ, mutation rate per learning event; ν, variance of demonstrator/tutor bias; p_att_, proportion of sllable types that are attractive to learn; α, conformity bias; and N_t_, number of potential demonstrators. Posterior inference was performed using Population Monte Carlo ABC (PMC-ABC). The authors computed 13 summary statistics per population, applied partial least squares (PLS) regression for dimension reduction, and then ran PMC-ABC on the resulting PLS features with decreasing acceptance threshold.

Using the published simulations output, we constructed population-specific posteriors by selecting, for each population, the 100 simulations most similar to the empirical PLS features (Euclidean distance) and fitting a kernel density estimator (KDE) to corresponding accepted parameter values. We then combined the six KDE-based posteriors into a collective posterior distribution using our method. For comparison, we also reconstructed the single posterior corresponding to conditioning on the mean empirical PLS feature vector across populations, as in Lachlan et al. Conditioning on the mean feature vector is a common ABC strategy for multi-replicate data, but it removes information about between-replicate variation.

#### SBI benchmarks

We also test our method on two common simulation-based inference benchmarks (44): *Gaussian Linear Uniform* (GLU), where the prior θ∼U[-1,1]^10^ and data points are drawn from a multivariate normal distribution such that y∼MVN(θ, Σ=0.1⊙I); and *Simple Likelihood Complex Posterior* (SLCP), which presents a more challenging simulation-based inference task. The prior is θ∼U[-3,3]^5^, and the observations are sets of four two-dimensional points sampled from a normal distribution whose mean and variance are nonlinear functions of θ. The resulting posterior has four symmetrical modalities and vertical cut-offs, making parameter inference particularly challenging.

For all tasks, we focused on n=10 replicates per observation set, aligning with realistic experimental limitations in biological settings (5,36,37,39,40).

#### Evaluation metrics

We assess the estimation accuracy of the inferred posterior by computing the log-ratio of the posterior mean and the true parameter value. To evaluate estimation uncertainty, we use *coverage* (45), defined as the probability that the true parameter is within an HDR of level α, P(θ_true_∈HDR_α_), for α = 0.1, 0.2,…, 0.9, 0.95. A perfectly calibrated posterior would obtain P(θ_true_∈HDR_α_)=α. While recent studies focus on the local classifier 2-sample test (46), we believe that the marginal metrics of accuracy and coverage better reflect the computational biologist’s needs from SBI. First, biological research is typically parameter-centric. Marginal metrics allow us to disentangle performance across distinct biological processes, verifying that key parameters are estimated reliably even if high-order correlations are imperfect. Global metrics like LC2ST compress this complexity into a single score, obscuring the specific sources of error. Second, calculating LC2ST requires training a separate classifier for every test observation, which becomes computationally prohibitive when validating performance across hundreds or thousands of synthetic multi-dimensional observations. In contrast, marginal accuracy and coverage can be efficiently computed, enabling us to perform high-throughput validation.

#### Hyperparameter tuning

Our implementation of the robust collective posterior distribution with SIR sampling includes a single hyperparameter: ε, the lower bound density of the robust individual posterior distribution, p_ε_(θ|x_i_). Based on our experiments, as long as the simulator and inference method do not change, the same value of ε remains useful, i.e., leads to both accurate inference and robustness to outliers. To avoid arbitrary selection, we derived a global and adaptive data-driven heuristics based on the dynamic range of the neural density estimator. First, we defined the baseline uninformative posterior density by drawing independent pairs of parameters and observations (θ,x) from the prior, p(θ), and the simulator, p(x|θ), respectively, and computing their conditional probabilities, p(θ|x), with the neural density estimator. A high (e.g., 95th) percentile of these probabilities represents the highest density the estimator assigns to a random pair (θ,x). Setting ε to the chosen percentile imposes a lower bound on the posterior density, ensuring that outlier observations are assigned a minimal weight in the collective posterior. This prevents outliers from disproportionately collapsing the collective posterior. We applied this approach to the synthetic benchmarks presented in the Results section. Second, in empirical applications, measurement noise and model misspecification (the *simulation-to-reality gap*) often cause the estimator to assign systematically lower posterior densities to empirical data compared to synthetic training data. To account for this shift, we propose an alternative estimation method. Here, ε is calculated specifically for each replicate set {x_i_}_i_ by evaluating the posterior densities of a batch of parameters {θ_j_}_j_ sampled from the prior, p(θ), conditioned on the replicates, {p(θ_j_|x_i_)}_i,j_. ε is then set to a chosen percentile of these posterior densities. This calibrates the minimum posterior density to the specific experimental conditions, ensuring robustness even when the overall posterior densities are shifted due to model misspecification.

#### Data and code availability

We use the Python package *sbi* (Tejero-Cantero et al. 2020) for rejection-ABC (MCABC in *sbi*), Neural Posterior Estimation (NPE), inference from multiple replicates with permutation-invariant embedding (NPE+PIE), and neural posterior score estimation (NPSE). We implemented the *collective posterior* in Python with PyTorch. All source code is available on GitHub at https://github.com/nadavbennun1/collective_posterior. Empirical data is publicly available from previous studies (5,36).

## 3. Results

### Synthetic data from evolutionary simulations

We evaluated the collective posterior and NPE+PIE (an established approach for inference from multiple replicates (30)) on the synthetic benchmark to assess posterior accuracy and calibration. The collective posterior achieved substantially improved empirical coverage across confidence levels (Figure 2d): coverage closely followed the nominal level, indicating well-calibrated uncertainty estimates, whereas NPE + PIE showed systematic under-coverage, consistent with overconfident posteriors (credible intervals that are too narrow). Across parameters (Figure 2e–j), the collective posterior yielded errors that were more tighly concentrated around zero, indicating higher accuracy and tighter posteriors around the true parameter values. Overall, the collective posterior produced more accurate and better-calibrated posterior estimates than NPE+PIE on this benchmark.

### Empirical data from evolutionary experiments

We next analyzed four empirical datasets from Chuong et al. (2025). Each dataset contains 5-8 experimental replicates of the same genotype (wildtype or one of three mutants). We generated 30,000 noisy Wright-Fisher simulations for training, adding Gaussian noise N(0,0.02) to the simulated time series. Using NPE as the replicate-level posterior estimator, the collective posterior produced accurate posterior predictive timeseries across datasets (Figure 3). In contrast, NPE+PIE yielded inaccurate posterior predictive timeseries for two of the four datasets (Figure 3ab). NPE training was also 10-fold faster than NPE+PIE with n_reps_=10 (Table S1).

**Figure 3.**
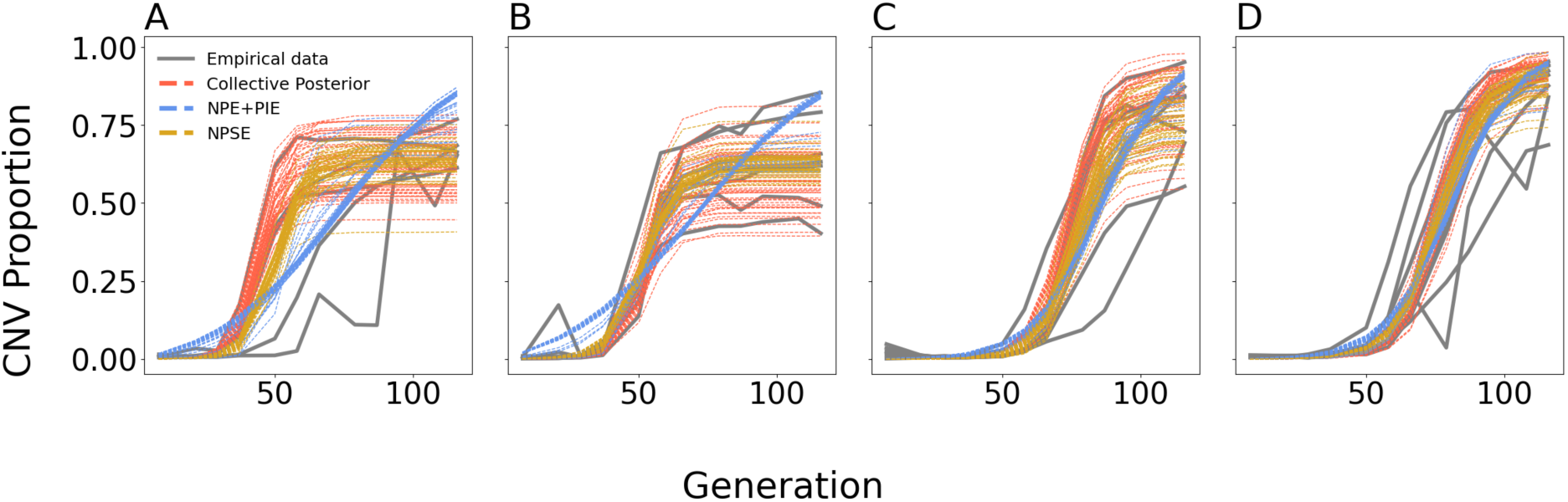
The collective posterior best matches the between-replicate variation in empirical evolutionary data. For each of four datasets from Chuong et al. (2025), we simulate 50 Wright-Fisher time series from the parameter sets from the collective posterior with NPE (red), NPE+PIE (blue), and NPSE (yellow), and compare them to empirical replicates (grey). The collective posterior more closely captures the observed between-replicate variability. Here, ε is estimated separately for each replicate set.

We additionally report NPSE results as an alternative approach to inference from multiple replicates. On these empirical datasets, NPSE generates posterior predictions that resemble plausible experimental replicates, but its aggregated posterior appears under-dispersed relative to replicate-to-replicate variability (see SI for an extensive comparison).

Furthermore, applying collective posterior inference with rejection-ABC (rather than NPE) as the replicate-level posterior estimator to the most challenging dataset (Figure 3b) produced MAP estimates similar to those obtained with NPE (Figure S2), providing an additional cross-check of our method.

### Robustness to outliers

Empirical replicate datasets often show greater dispersion than simulations, and replicates that pass standard outlier checks still induce anomalous inference. In its default form (eq. 1), the collective posterior can be influenced by such outlier replicates. To therefore use a robust posterior formulation that enforces a minimum per-replicate posterior density ε (eq. 2). Figure 4 illustrates the effect on one dataset from Chuong et al. (2025): with an extremely small floor (ε=e^-1000^), the collective posterior is dominated by a single outlier replicate. Increasing the floor to ε=e^-10^ yields a collective posterior that aligns with the high-density overlap of the replicate-specific posteriors and substantially improves posterior predictive checks (Figure 4).

**Figure 4.**
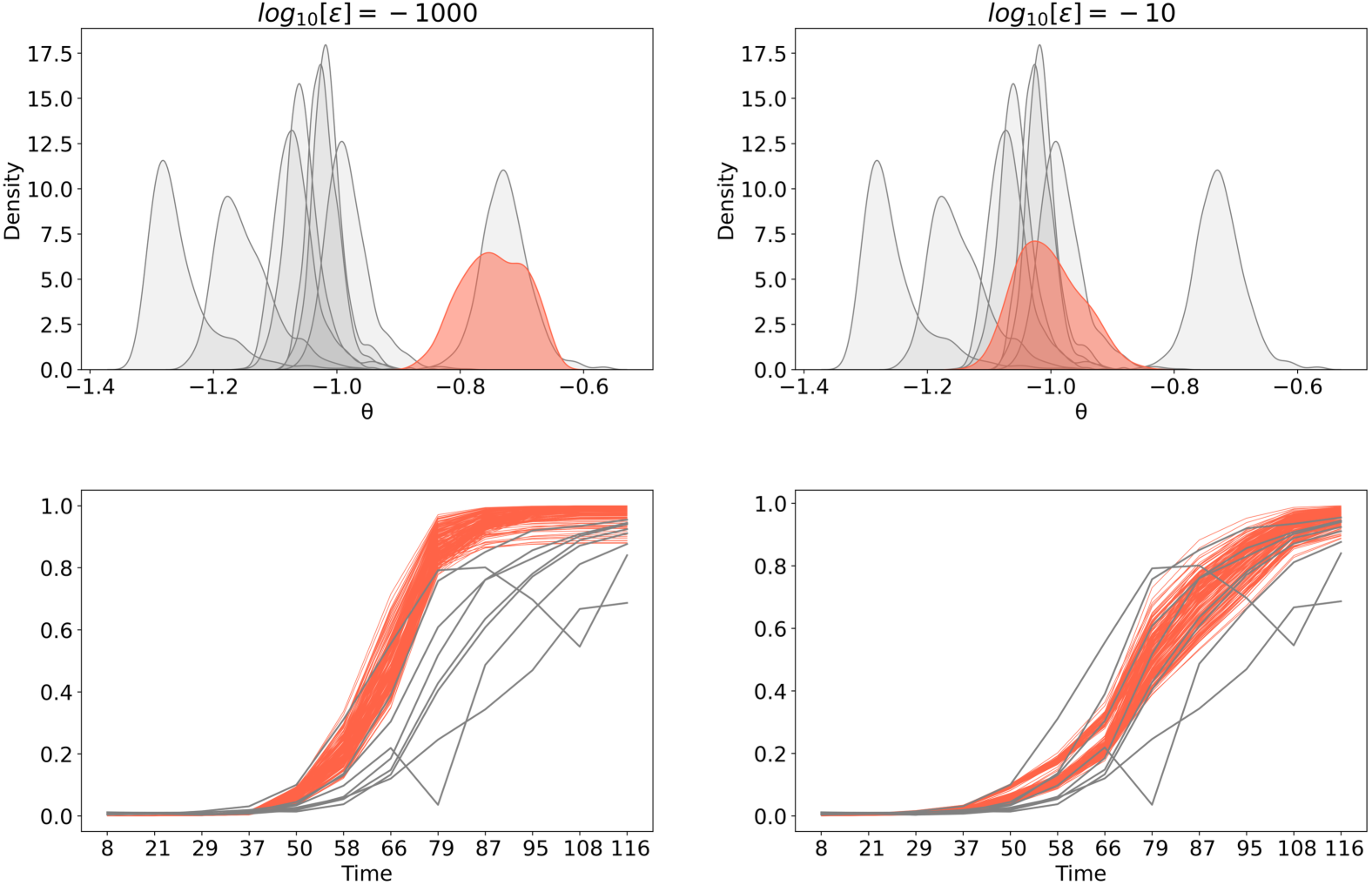
Robust collective posterior inference using the ε floor. **(A-B)** Inferred posteriors for the dataset in Figure 3D under two choices of the minimum-density parameter ε. With log(ε)=-1000, the collective posterior (red) is dominated by an outlying replicate; with log(ε)=-10, the collective posterior (red) shifts toward the region of overlap among replicate-specific posteriors (grey). **(C-D)** Posterior predictive checks: empirical replicate time series (grey) and 200 simulations from parameter draws from the collective posteriors (red), showing improved agreement with the between-replicate variability under the higher ε setting.

### Swamp sparrow song dataset

We analyzed the swamp sparrow song dataset of Lachlan et al. (5), which comprises six geographically distinct populations treated as replicates. The goal is to infer five parameters of a cultural transmission model from the population-level summary statistics derived from recorded song repertoires, and to compare posterior inference based on combining replicate-specific evidence versus conditioning on summary statistics averaged across populations.

Using the published simulation output from Lachlan et al., we reconstructed replicate-specific posteriors for each population by smoothing the accepted parameter sets with kernel density estimation. These replicate-specific posteriors were more concentrated and yielded lower discrepancies to the empirical summary statistics than the posterior obtained by conditioning on mean summary features across populations (Figure 5).

**Figure 5.**
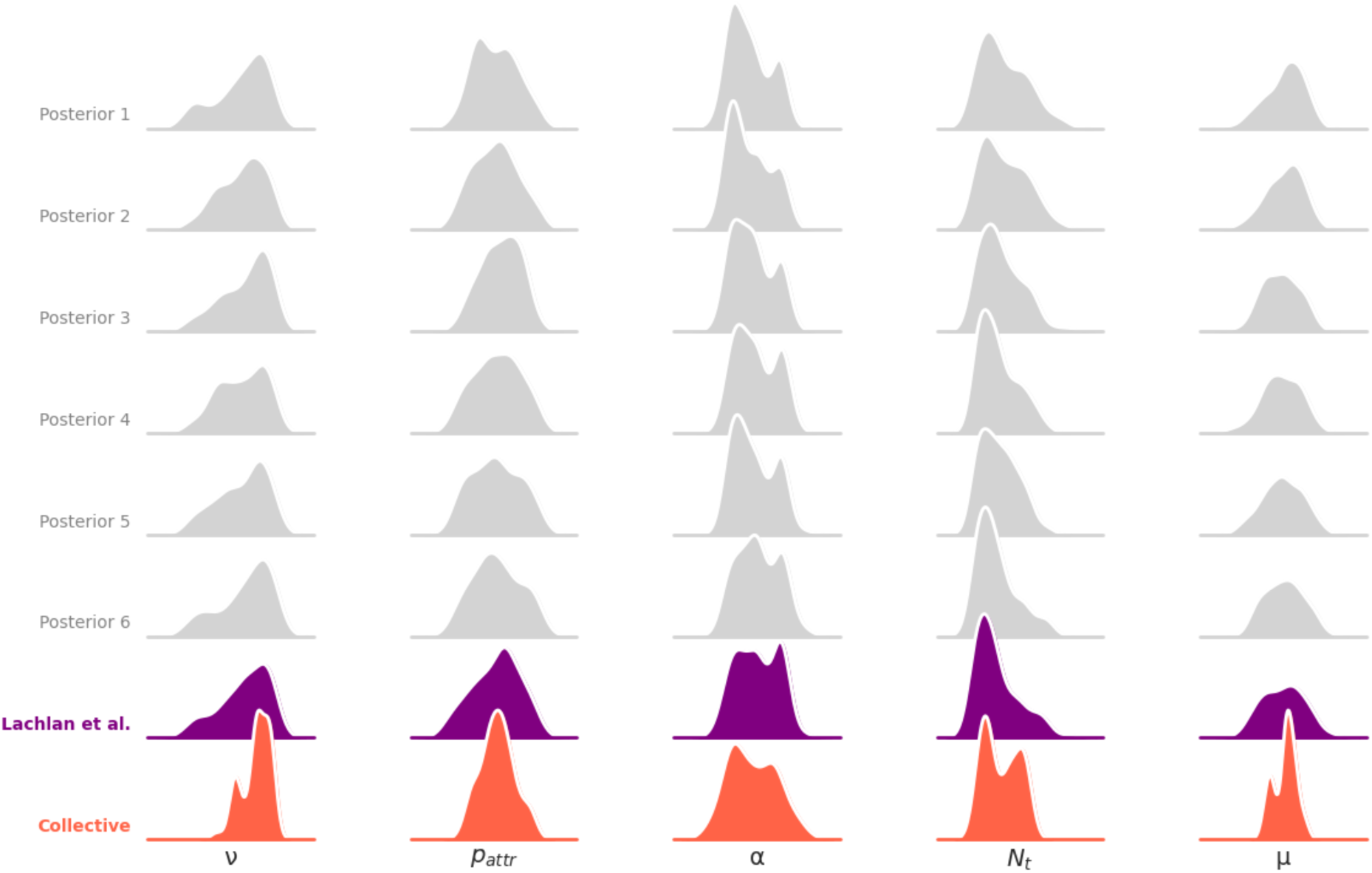
The collective posterior infers cross-population consensus in swamp sparrow cultural evolution parameters. Grey curves show replicate-specific posteriors inferred separately for six populations. The purple curve reproduces the posterior from Lachlan et al. (2018), obtained by conditioning on summary statistics averaged across populations, which yields a comparatively broad posterior. The collective posterior (red) combines replicate-specific posteriors to emphasize parameter regions supported consistently across populations, producing a sharper posterior for parameters with strong cross-population agreement (e.g., ν and μ).

We then combined the six replicate-specific posteriors using the collective posterior. It assigned high density to regions of parameter space where the replicate-specific posteriors overlap, and low density where they disagree (Figure 5). In contrast, the mean-conditioned posterior of Lachlan et al. remains comparatively broad, assigning high density over a wider range and therefore more closely resembling the replicate-specific posteriors. The increased concentration of the collective posterior is analogous to a simple Gaussian product-of-experts case: the product of *k* identical Gaussian distributions N(μ,σ^2^) is proportional to a Gaussian with the same mean and reduced variance, 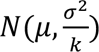. Our implementation exhibits this variance reduction, reflecting the degree of agreement among replicates. More generally, these results show that the collective posterior captures cross-replicate consensus by combining replicate-specific posteriors, rather than by first averaging or their summary statistics and then performing inference on the aggregated data.

### SBI benchmarks

We next evaluated the collective posterior on two standard SBI benchmarks (GLU and SLCP), using NPE as the replicate-level posterior estimator, and compared it to NPE+PIE. Across both benchmarks, the collective posterior was comparable to NPE+PIE, and in some cases, more closely centered on the true parameter values. For GLU, where averaging replicates is informative because the mean of the replicates concentrates around the truth, the collective posterior achieved higher accuracy and better calibration (Figure S8). For SLCP, performance was similar overall, with improved accuracy for one parameter (Figure S9). Together, these benchmark results are consistent with the patterns observed in the empirical datasets.

## 4. Discussion

Taken together, our results show that collective posterior inference is more accurate and better calibrated than NPE+PIE across tasks we considered, including three synthetic benchmarks and one empirical dataset. In our analyses, it required significantly less training than NPE+PIE while remaining compatible with a wide range of replicate-level posterior estimators.

Based on our findings, as well as related results in the literature (32–35), we conclude that combining replicate-specific posteriors can be advantageous relative to directly learning a single multi-replicate posterior, particularly when replicate-to-replicate variability and departures from the assumed generative model are non-negligible. A central difficulty in inference from multiple replicates is accommodating variability that exceeds what is expected under the model’s intrinsic stochasticity. Such excess between-replicate variance can arise from unmodeled hierarchical structure, technical effects, or other sources of heterogeneity not captured by the simulator. We show that, even in the presence of such excess variance, collective posterior inference can still yield accurate and well-calibrated posteriors.

An alternative strategy is to explicitly model the full hierarchical data-generating process and concatenate the replicates into a single “super observation” vector, followed by inference on the global parameters from this high-dimensional input. However, posterior estimation generally becomes harder as the dimensionality of the data increases (curse of dimensionaility), both for neural and non-neural density estimators (26,44,47). Consistent with this, Avecilla et al. (2022) found that concatenating replicates reduced inference accuracy when estimating selection coefficients from evolutionary experiments.

The collective inference approach is method-agnostic: it can be combined with any replicate-level posterior estimator to perform inference from multiple replicates. A key feature is its robustness to outliers: during aggregation, the method can downweight or effectively ignore idiosyncratic replicates without requiring explicit outlier detection (Figure 4). On synthetic benchmarks, the collective posterior paired with NPE matched or outperformed NPE+PIE, even though both approaches use the same NPE posterior estimator and differ primarily in how replicate information is incorporated (aggregation of replicate-level posteriors versus set-conditions via premutation-invariant embeddings.)

We showed that collective posterior inference can accommodate biological variability and idiosyncratic replicates while preserving accurate parameter estimation. Because it operates by combining replicate-level posteriors, it is compatible with a broad range of likelihood-based and likelihood-free posterior estimators, including newer neural SBI methods. Moreover, unlike set-conditioning approaches that assume a fixed maximum number of replicates, the collective posterior naturally scales to varying numbers of replicates.

Finally, the collective posterior can also be applied to *computational* replicates, such as multiple independently trained neural posterior estimators, providing a “floor raising” alternative to standard ensembling. Whereas common ensembles (e.g., bagging or simple averaging) combine estimators by averaging their densities, yielding a broad union that retains between-run training variability (18, blue), our robust product formulation (Eq. 7) aggregates estimators through an intersection-type operation that emphasizes regions where they agree. As a result, it attenuates run-to-run training variance and produces sharper posterior estimates that more directly reflect the consensus signal (Figure S18, red).

## Authors’ Statement

We used ChatGPT 5 and Gemini 3 for text editing and code optimizations.

## Acknowledgements

We thank all members of the Ram lab, Julie N. Chuong, Titir De, Yoni Green, Tal Pupko, Adi Stern, and Ofir Levy for helpful discussions and advice. This work was supported in part by the US–Israel Binational Science Foundation (YR & DG 2021276), Minerva Center for Live Emulation of Evolution in the Lab (YR), and fellowships from the Edmond J. Safra Center for Bioinformatics at Tel Aviv University (NBN) and the AI and Data Science Center at Tel Aviv University (NBN).

## Supplementary Information

### Simulation-based Inference

**Figure S1.**
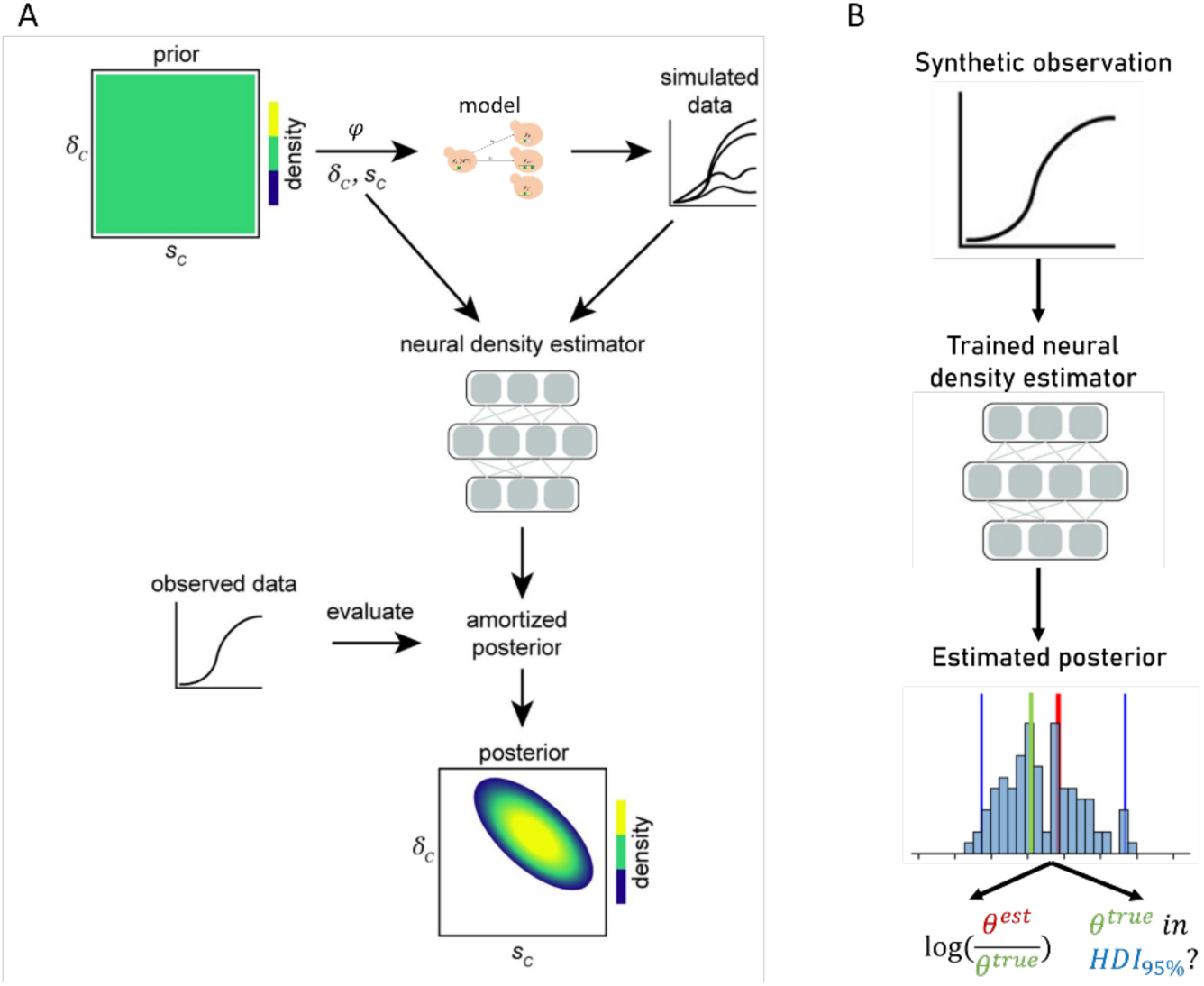
Simulation-based inference with Neural Density Estimation. **(A) Inference.** Proposal parameters are sampled from a prior distribution and used for model simulations. These parameters and the corresponding simulation results are used as a training set for a neural density estimator, e.g., normalizing flow. The neural density estimator evaluates an observation, effectively approximating a joint posterior distribution of model parameters. Adapted from Cranmer, Brehmer, and Louppe, 2020 (1). **(B) Evaluation.** The trained neural density estimator is evaluated on synthetic observations for which the true parameters are known. By comparing the resulting posterior distributions to the true parameters, we estimate the accuracy of the sample median (or another point estimate) and the coverage of the highest-density region (HDR).

### Full Derivation of the Collective Posterior Distribution

Let *x*_1_,…, *x*_*n*_ be empirical observations such that *x*_*i*_∼*p*(*x*|*θ*). Assuming we have already estimated *p*(*θ*|*x*_*i*_) using NPE, we wish to estimate *p*(*θ*|*x*_1_,…, *x*_*n*_). Using Bayes’ theorem,

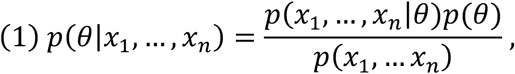

Assuming *x*_*i*_, *x*_j_ are independent given *θ*, i.e., *p*(*x*_*i*_, *x*_j_T*θ*) = *p*(*x*_*i*_|*θ*)*p*(*x*_j_T*θ*), we get

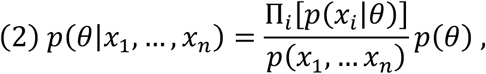

Using Bayes’ rule again,

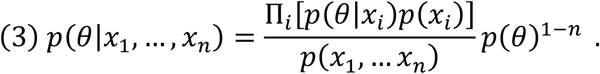

Next, recalling the law of total probability

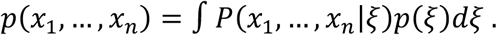

Using Bayes’ rule,

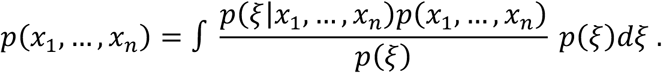

As *p*(*x*_1_,…, *x*_*n*_) is positive and independent of *ξ*, we can taKe it out of the integral and cancel it out from both sides,

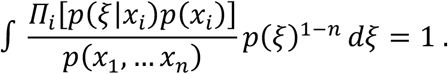

Using Bayes’ rule in (4),

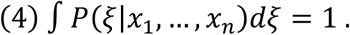

Using commutativity of multiplication,

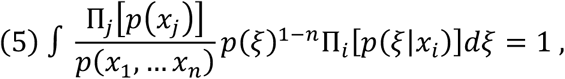

Where *p*(*ξ*), the prior, is Known to us, and *p*(*ξ*|*x*_*i*_), the individual posteriors, are estimated by a density estimator. Therefore, only the marginal liKelihoods *p*(*x*_*i*_) and *p*(*x*_1_,… *x*_*n*_) are unKnown, yet positive and independent of *ξ*.

We can therefore consider 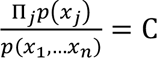 and get

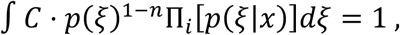

Then, plugging C in (3) will result in the collective posterior distribution as in eq. 1,

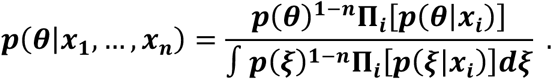

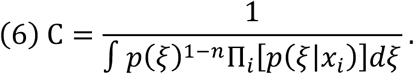

### Additional Sampling Strategies

#### Markov Chain Monte Carlo (MCMC)

As a robust alternative for exploring complex local landscapes, we implemented a Metropolis–Hastings sampler initialized from high-probability modes. We first generated a large pool of candidates from the prior and evaluated their unnormalized posterior probabilities. The 50 candidates with the highest log-probabilities were selected to initialize 50 independent Markov chains. Each chain proceeded with Metropolis– Hastings updates to locally explore the posterior mass, ensuring adequate coverage of the collective posterior’s peaks even in cases where the global proposal distribution of SIR might be inefficient.

#### Rejection Sampling

We also implemented exact rejection sampling. This required first approximating the normalizing constant 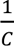 (the denominator of Eq. 1) via Monte Carlo integration using *N*_*c*_ = 10^5^ prior samples, and averaged across 10 independent approximations. Once the constant was estimated, samples were drawn from the prior and accepted or rejected based on the ratio of the collective posterior probability to the proposal. While asymptotically exact, this method proved computationally prohibitive for high-dimensional or highly peaked posteriors due to vanishing acceptance rates.

#### Hyperparameter tuning

Sampling with MCMC or rejection sampling introduces two additional hyperparameters: *N*_*samples*_, the number of initial candidate samples to keep for MCMC sampling; and *N*_*c*_, the number of prior samples to evaluate the denominator of eq. (1) for sampling from the posterior distributions.. For *N*_samples_, we set a default of 50, corresponding to the 50 candidate points with the highest log-probability under the estimated normalization constant 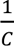. These candidates serve as initialization points for independent MCMC chains, ensuring sufficient exploration of multiple posterior modes while avoiding over-concentration in a single region of high probability. For comparison, *sbi* typically employs 20 independent MCMC chains, following the recommendations of Neural Likelihood Estimation (2). We extended this number since a collective posterior distribution can potentially have a larger number of modalities, therefore requiring a broader inspection of the support.

Finally, sampling from the collective distribution requires a one-time computation of *C*, with computation time scaling linearly with *N*_*c*_, increasing from a few seconds for *N*_*c*_ = 10^5^ to about a minute with *N*_*c*_ = 10^6^ on a 48-core CPU.

### The Collective Posterior Distribution With ABC

**Figure S2.**
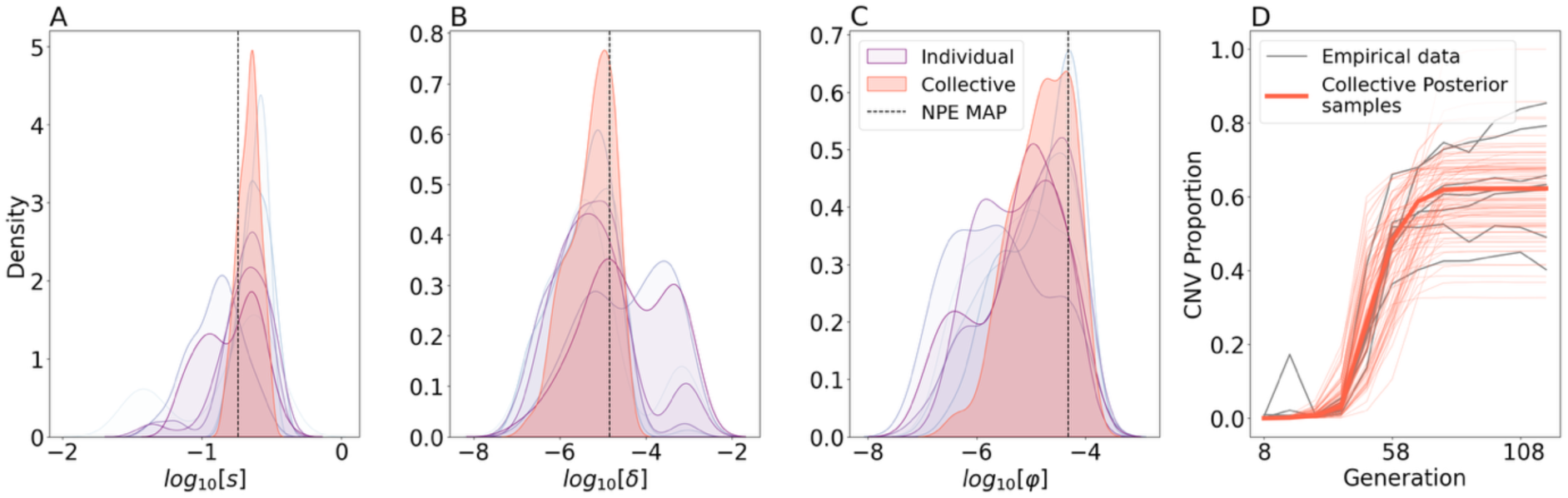
The collective posterior distribution is similar, regardless of the individual inference framework. **(A-C) The collective posterior distribution from ABC concentrates around the NPE Maximum A-Posteriori (MAP).** Individual posteriors of seven experimental replicates inferred for the Wright-Fisher model with rejection-ABC (purple, *sum of squared distance* ≤ 0.5, 10,000 simulations per observation) compared to the collective posterior distribution (red). Collective MAP estimates of NPE (3) are marked by the black dashed lines. **(D) Collective posterior predictions agree with empirical data.** Collective posterior predictions using Wright-Fisher simulations with 100 parameter sets sampled from the collective posterior distribution (thin red lines), from the mean of the collective posterior (bold red line), and empirical data (grey).

### Wright-Fisher model (3)

**Figure S3.**
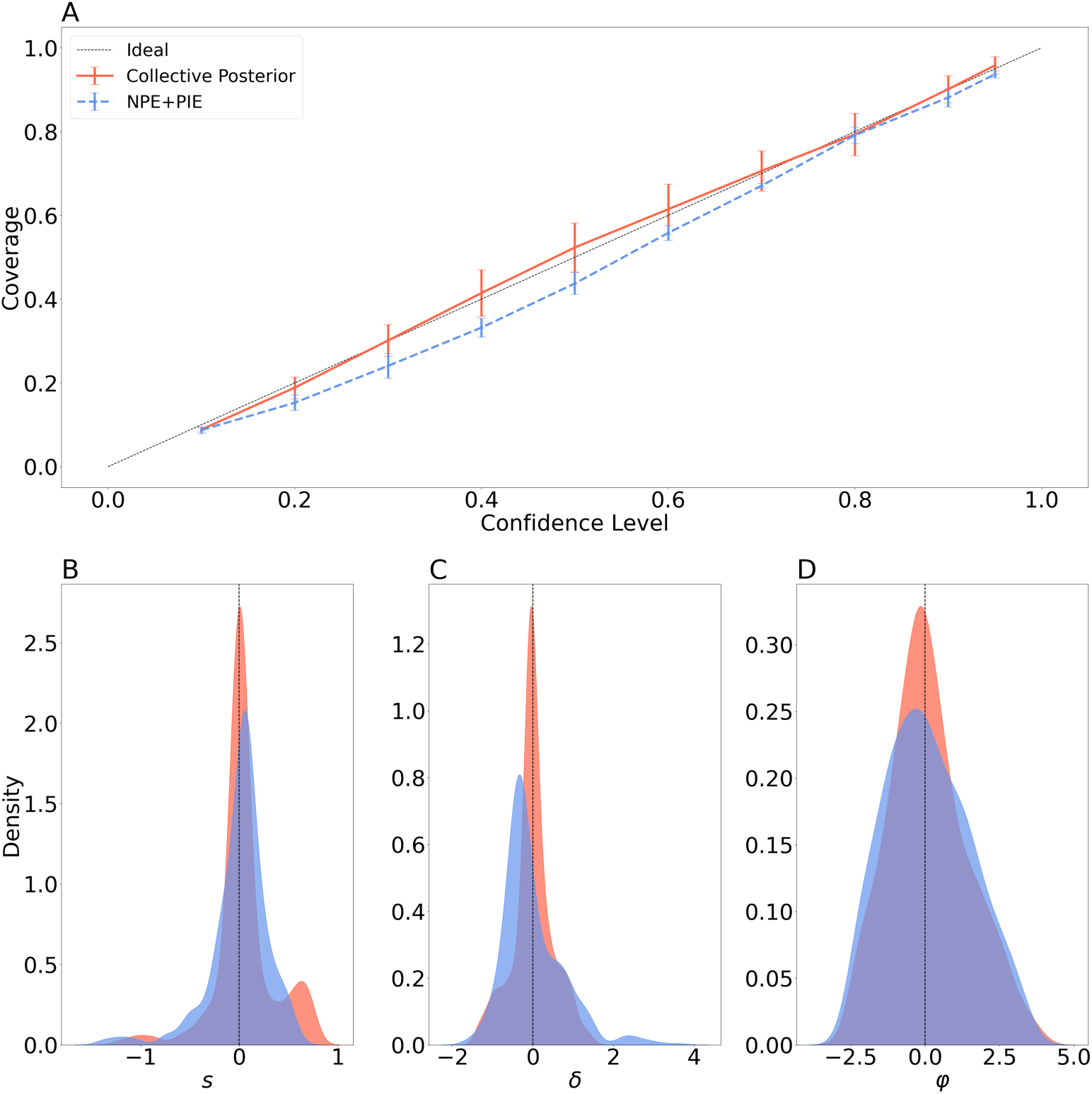
Collective posterior distributions from synthetic Wright-Fisher simulations. **(A-C). Accuracy.** The difference between the sample mean and the true parameter. The median errors for the collective posterior distribution are 0.0144, −0.0354, and −0.0550 for *s*, *δ*, *φ*, respectively. The median errors for NPE+PIE are 0.0492, −0.2277, and −0.0086, respectively. **(D) Coverage.** The probability that the true parameter is within the *HDR*_*α*_ of the estimated posterior, for *α* ∈ {0.1,0.2,…, 0.9,0.95}. Coverage is nearly identical for the two methods across all parameters and confidence levels, with a few minor exceptions.

**Figure S4.**
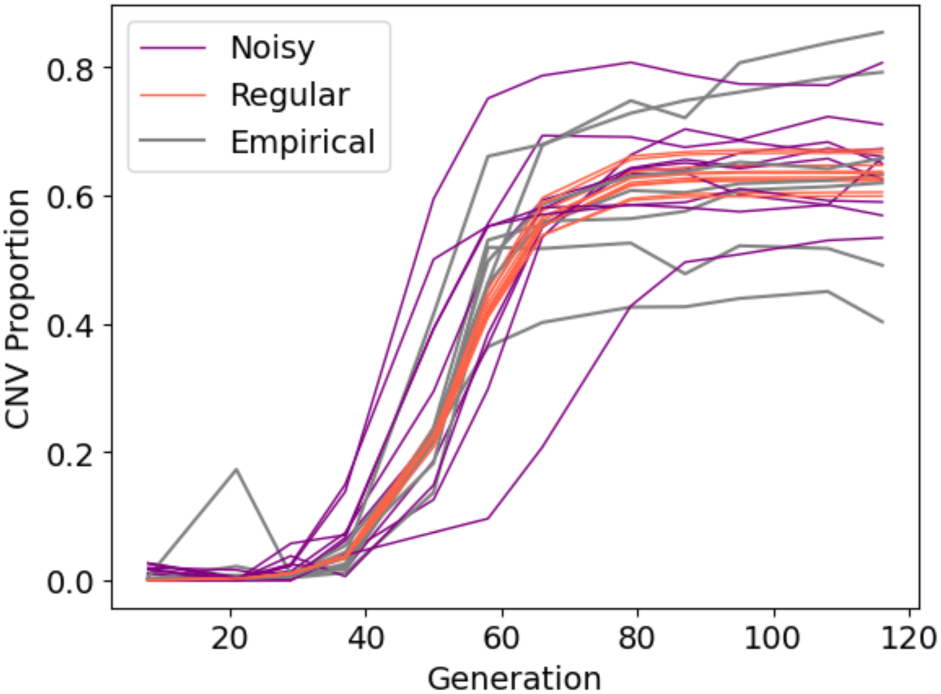
Our noising scheme can lead to experimental-like variability. Estimated parameters for the LTRΔ genotype from Chuong et al. (2025) were simulated ten times using the standard WF simulator (regular) and using perturbations on *θ* and Gaussian measurement noise. Empirical data is shown in grey.

**Figure S5.**
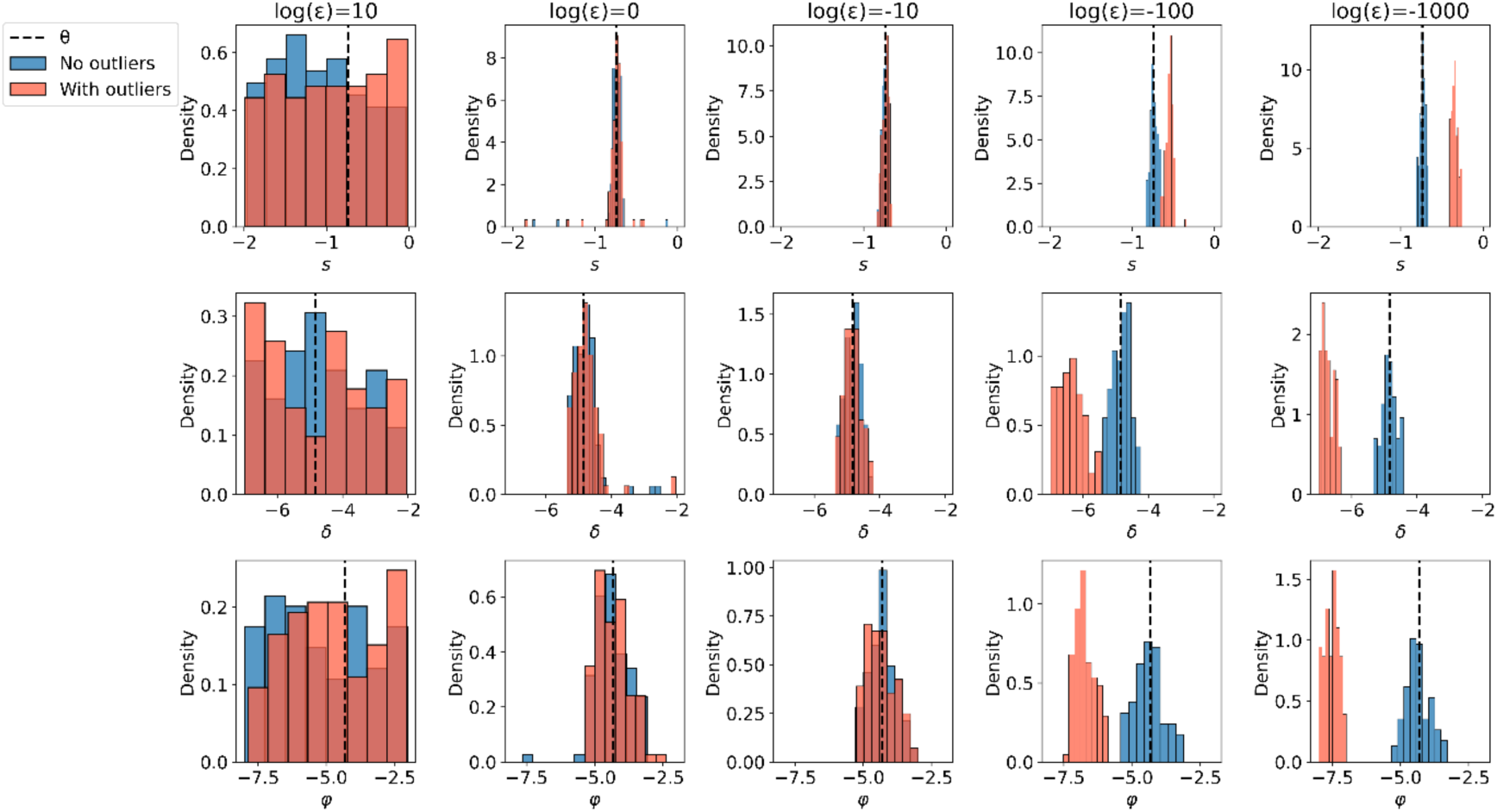
Effect of *ε* on inference. We used *θ* = log_10_(0.18, 1.44 ⋅ 10^−5^, 4.78 ⋅ 10^−5^) (one MAP estimate from Chuong et al., 2025) for simulating 10 observations. Then, we simulated 2 outliers using log_10_(*θ* + 0.25) and log_10_ (*θ* − 0.25). Each column shows the marginal posterior distributions for the three model parameters using a different value of *ε*, when inferring a collective posterior distribution from the 10 standard simulations (“No outliers”, blue) or the 12 mixed simulations (“With outliers”, red). Without outliers, inference is accurate for any *ε* ≤ 1. With outliers, however, the standard collective posterior distribution (*ε* → 0) fails to infer the parameters accurately. Our implementation of the collective posterior distribution exhibits accurate inference with robustness to outliers for *ε* = *e*^−10^.

### Score-based methods for SBI from multiple observations

Neural posterior score estimation and diffusion models (4–6) have shown promise for simulation-based inference from i.i.d replicates. We therefore compared NPSE with the Gaussian sampler (5) to collective posterior inference with standard NPE on both the synthetic and empirical data.

To train NPE, we used Masked Autoregressive Flow (MAF) (7) as the neural density estimator. MAF is recommended mostly for its training time, while considered subpar in terms of accuracy. Both Geffner et al. and Linhart et al. showed that their methods outperform NPE with MAF on several benchmarks, meaning that MAF is not the optimal replicate-specific posterior estimator. Here, we show that our collective posterior inference, with MAF as its neural density estimator, slightly outperforms NPSE in the noisy biological setting (Figure S6). Importantly, our method is orders of magnitude faster at inference, with similar training times (Table S1).

Arruda et al. (6) introduced a score-based method for efficiently estimating a posterior distribution from a large number of i.i.d. replicates and simulators with many parameters. While the method is promising, it requires significant hyperparameter tuning, which is non-trivial for general users, and is limited to a specific (score-based) density estimator. We recommend attempting their method, especially for massive datasets or parameter-heavy simulators. However, our collective posterior inference is agnostic to the density estimator and is essentially free of hyperparameter tuning, making it extremely applicable to new problems.

**Figure S6.**
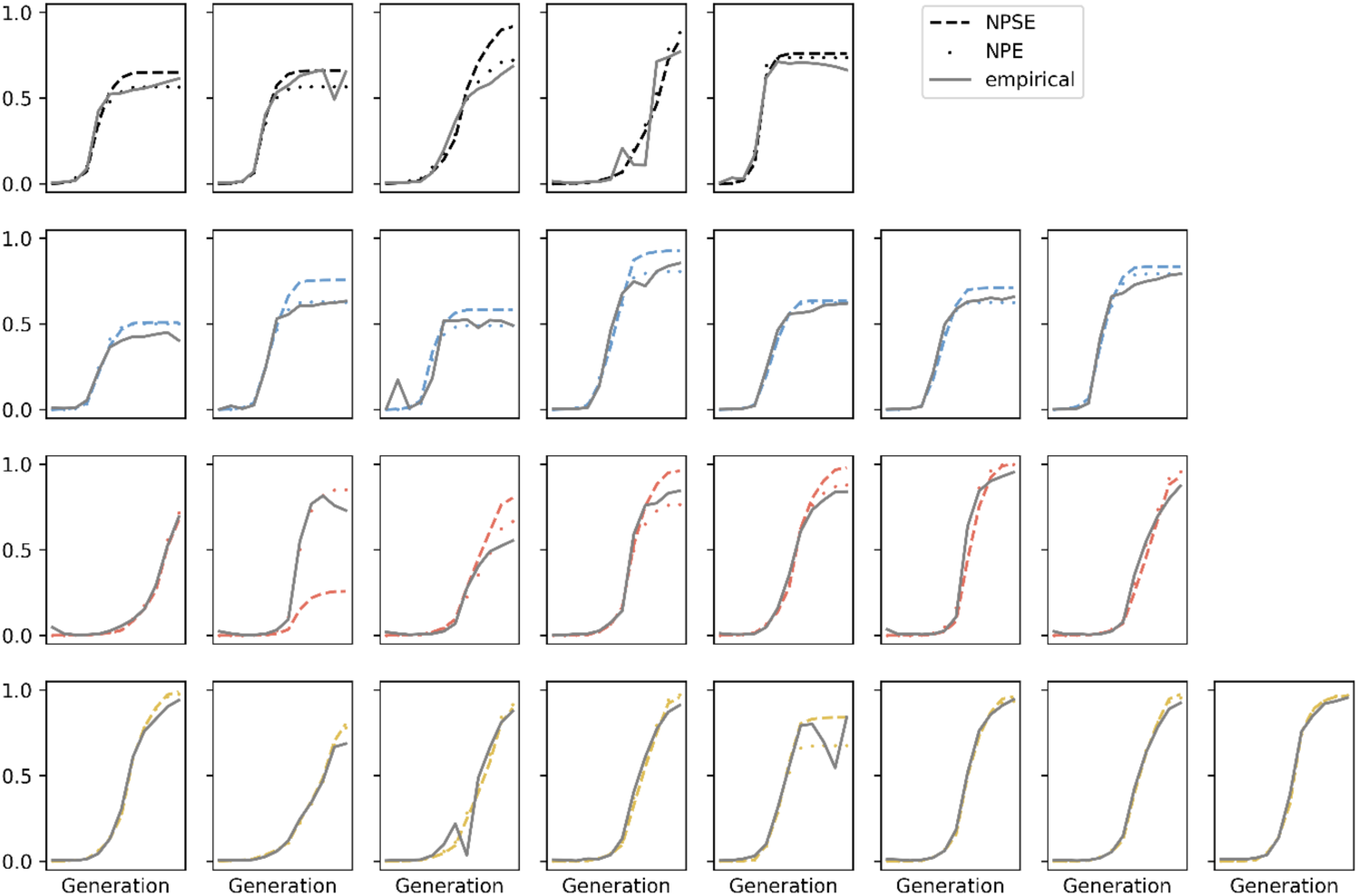
Posterior predictive checks of NPE and NPSE on the empirical data. Simulation of posterior sample mean from 100 samples. Both density estimators trained on 30,000 noisy WF simulations with their default hyperparameters.

**Figure S7.**
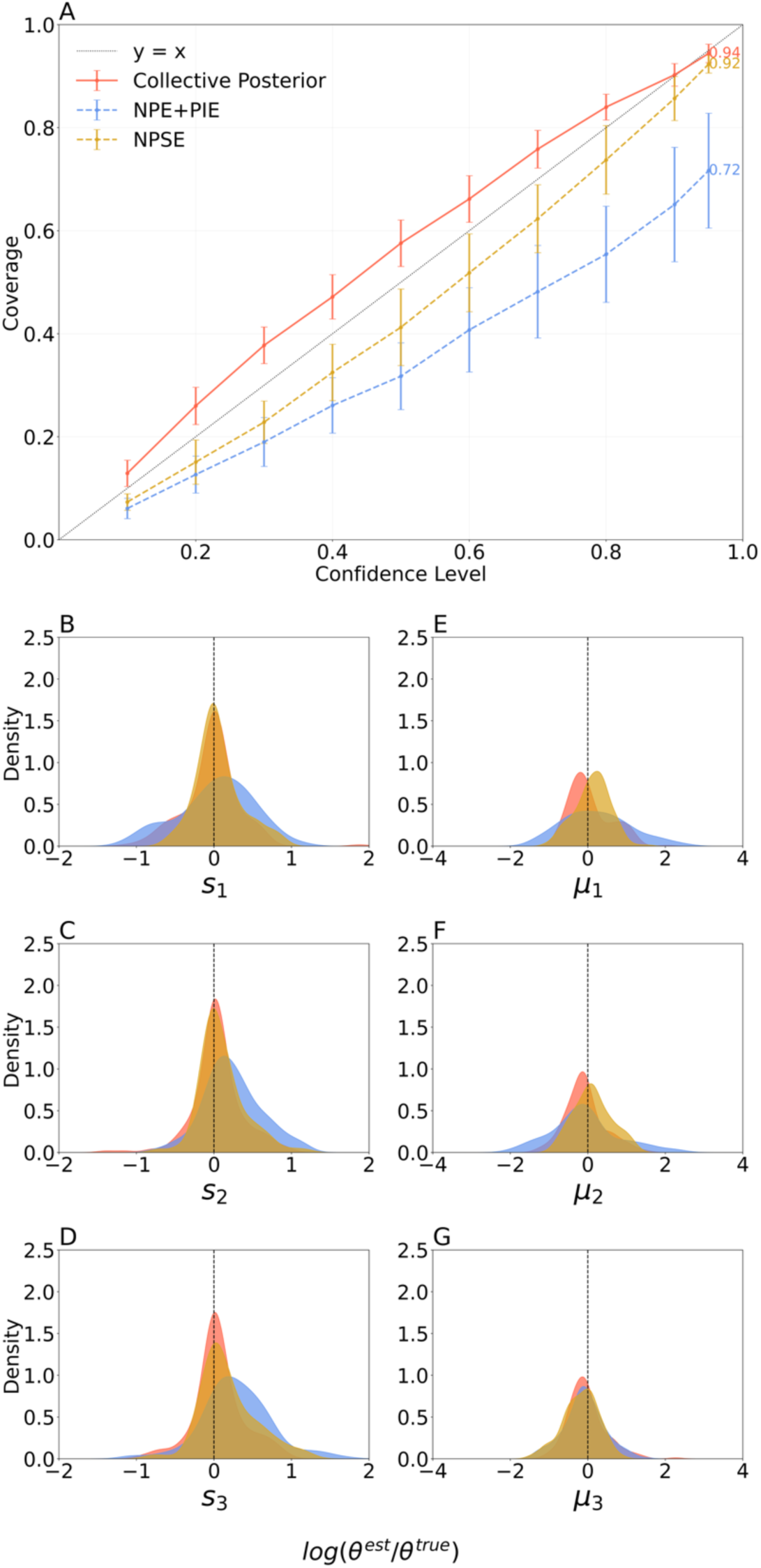
The collective posterior with NPE slightly outperforms NPSE on noisy evolutionary simulations. **(A) Coverage plots.** Box plots of coverages Pr[*θ*_*i*_ ∈ *HDR*_*α*_], corresponding to confidence levels *α*. The HDRs of the collective posterior consistently include the true parameter at higher probabilities. **(B-J) Centrality.** Distributions of the difference between the posterior sample median and the true parameter for all model parameters. The collective posterior has a higher density at zero for all parameters, indicating a better concentration around the true parameter.

### SBI Benchmarks

**Figure S8.**
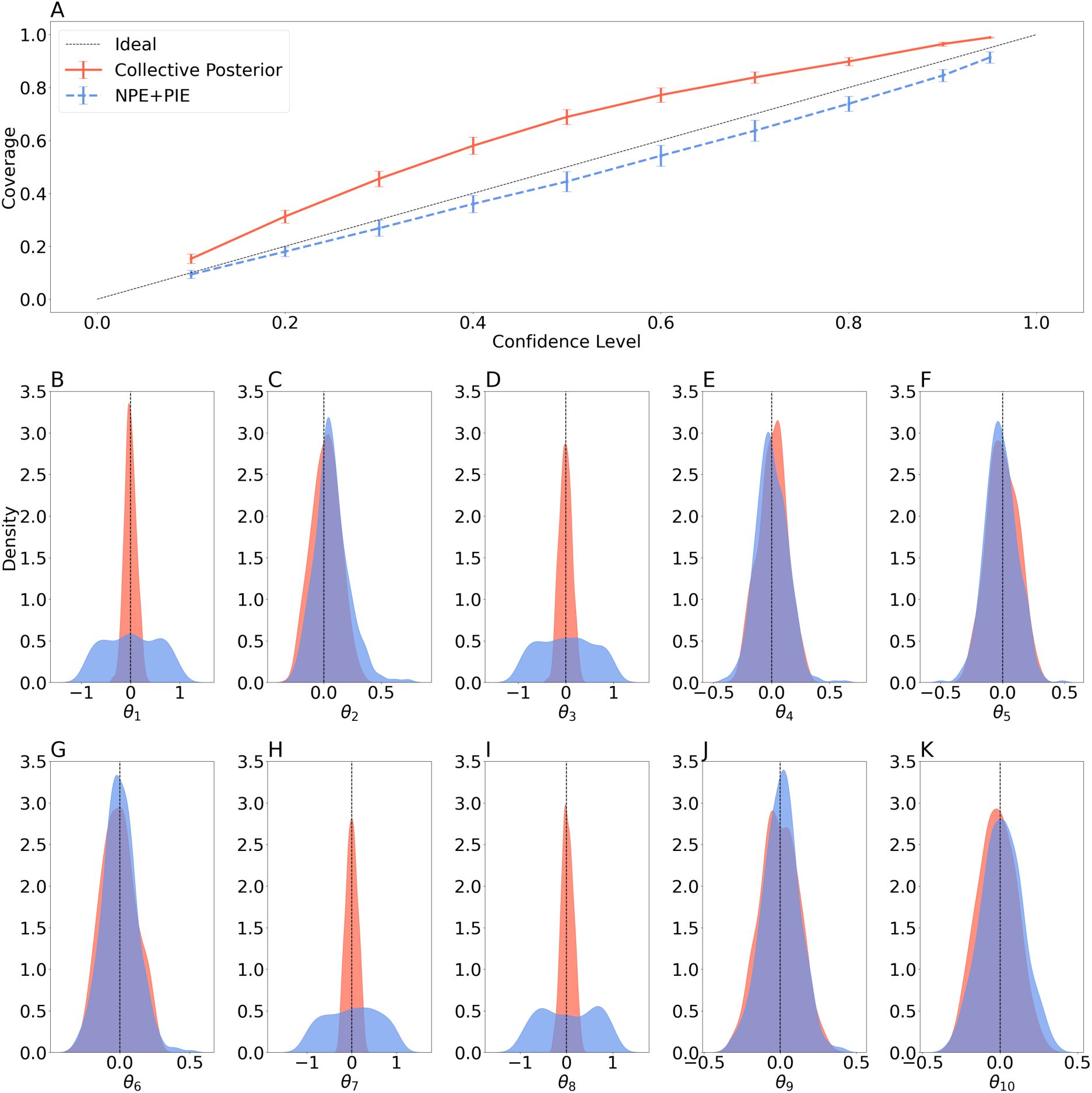
The sample median accuracy of the collective posterior and NPE + Permutation Invariant Embedding (PIE) for Gaussian Linear Uniform (GLU). Subplots show 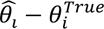 for the ten estimated parameters.

**Figure S9.**
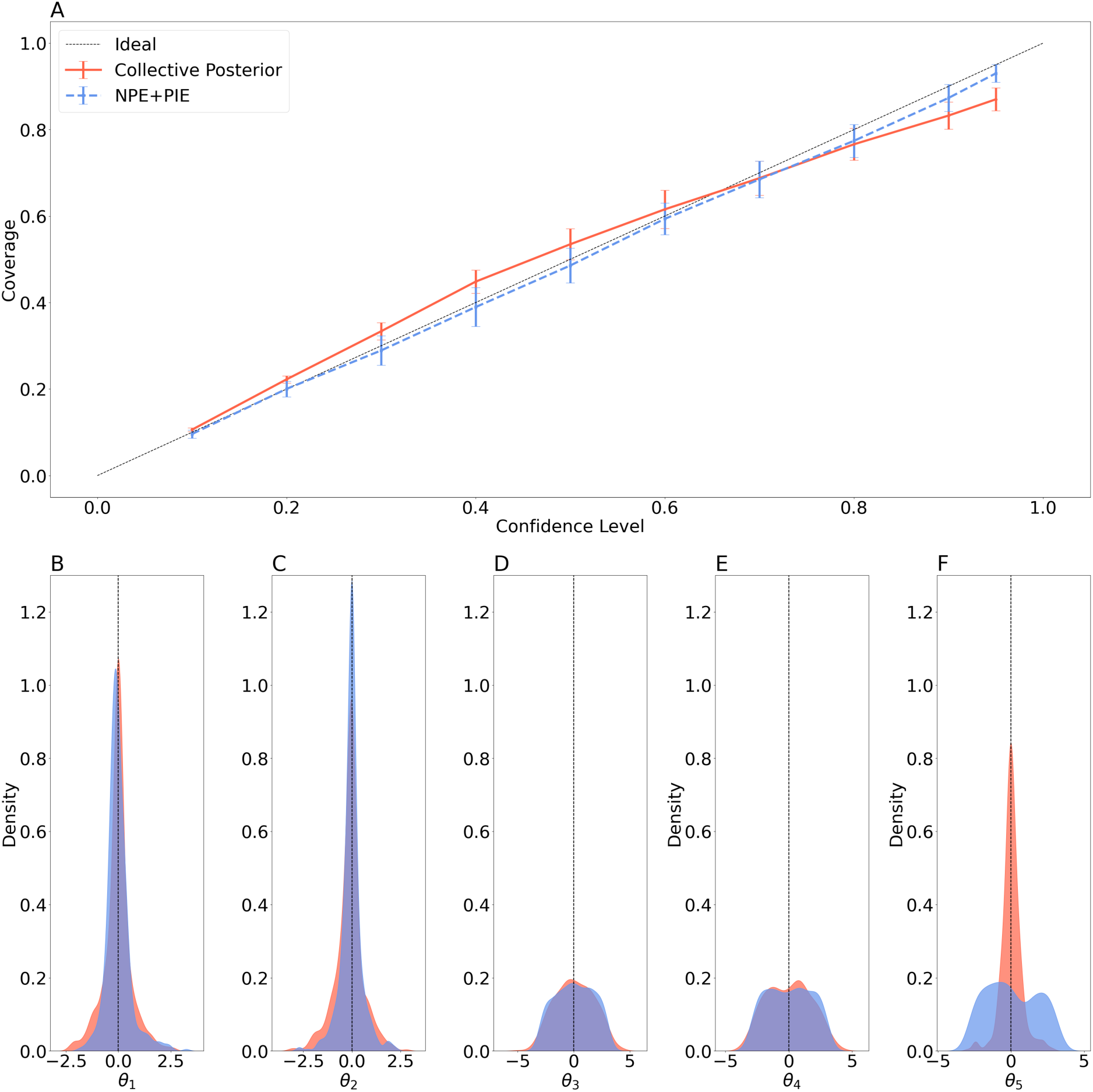
The sample median accuracy of the collective posterior (with rejection sampling) and NPE + Permutation Invariant Embedding (PIE, with rejection sampling) for Simple Likelihood Complex Posterior (SLCP). Subplots show 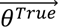 for the five estimated parameters.

**Figure S10.**
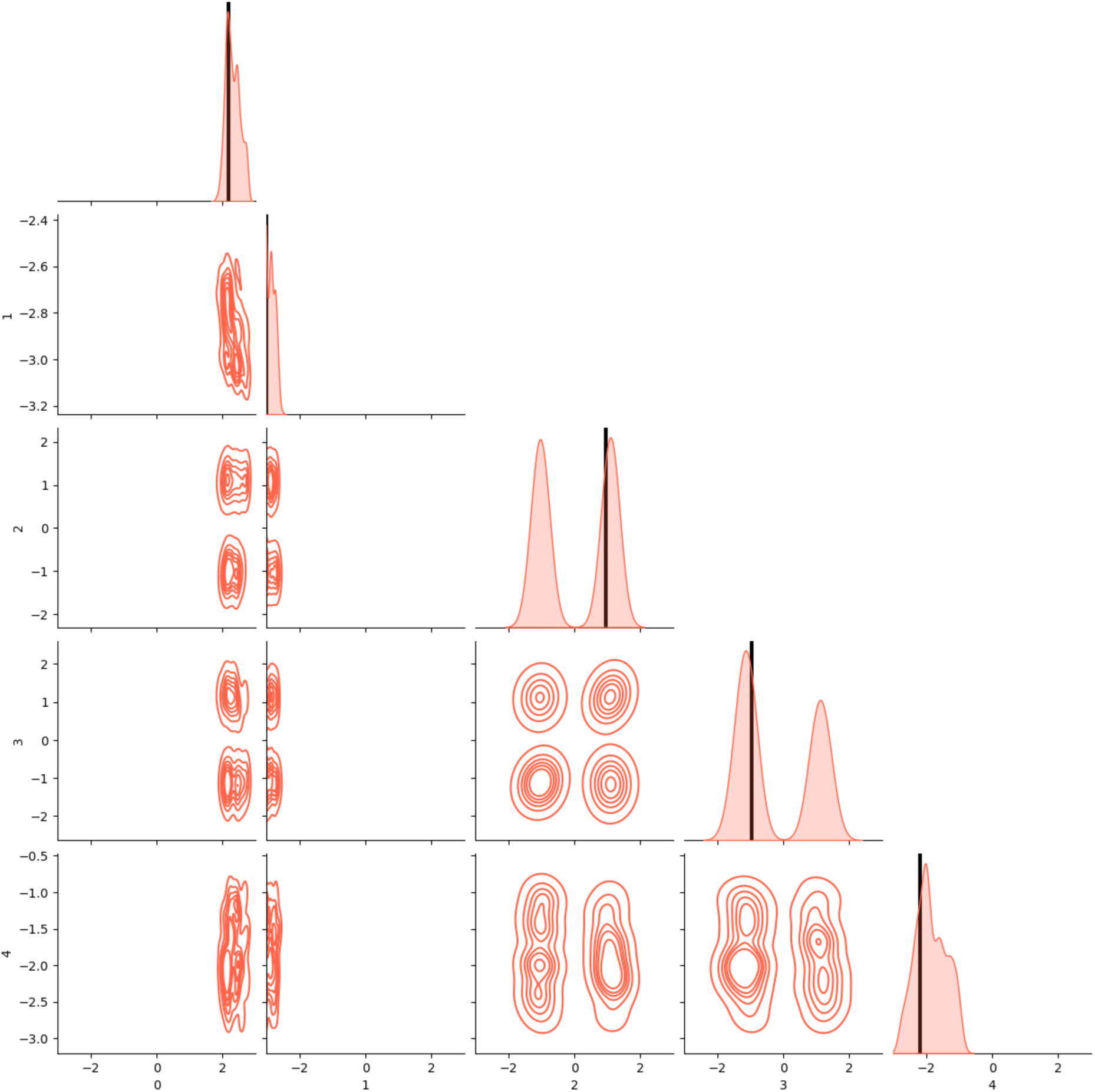
Example of pairwise and marginal collective posteriors for SLCP. A single collective posterior distribution for SLCP, with 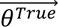 = (2.1796, −2.9841, 0.9612, −0.9603, −2.1850) (the last benchmark from (8)), marked in black vertical lines. The observation is a set of 10 independent SLCP simulations of 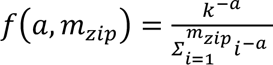.

### An Additional (unrealistic) Test Case – Genome Rearrangement

We applied our method to a recent genome rearrangement model (9), which models genome evolution along a phylogenetic tree due to rearrangement events: inversion, a reversal of a block of genes in its place, with a rate *R*_*in*_; translocation, a relocation of a block of genes, with a rate *R*_*tr*_; fusion, merging of two chromosomes to one, with a rate *R*_*fu*_; fission, splitting of one chromosome to two, with a rate *R*_*fi*_. Block sizes (for inversion and translocation) follow a trimmed Zipfian distribution with an estimated exponent parameter *a* and a fixed maximal value *m_zip_* = 20, such that 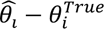. The model output is a set of genomes, one for each of the taxa on the terminal nodes of the phylogenetic tree. This model has high variance among simulations with the same parameter values, and we further increase this variance by considering a very large number of chromosomes in the ancestral genome, 90 (Figure S10). Moshe et. al validated their simulation-based inference method on synthetic data and applied it to a set of eighteen *Candida* species (10). Here, we simulated ten independent synthetic observations using their estimated parameters. Then, we estimated the individual posteriors using rejection-ABC and aggregated them to a collective posterior distribution.

We generated ten synthetic observations using the parameters estimated by Moshe et al. (2022). Using rejection-ABC inference for individual posterior distributions, the collective posterior distribution concentrated around the true parameters (Figure S11). Notably, the collective posterior had the highest density for the true value of all parameters and was bi-modal for *a* and *R*_*fu*_(Figure S11). This bimodality reflects the diverse individual posterior distributions. While the simulator applies to empirical data, it is highly unrealistic to obtain multiple independent observations with the same phylogenetic tree. We chose to include this example to show the applicability of the collective posterior to additional types of biological simulators.

**Figure S11.**
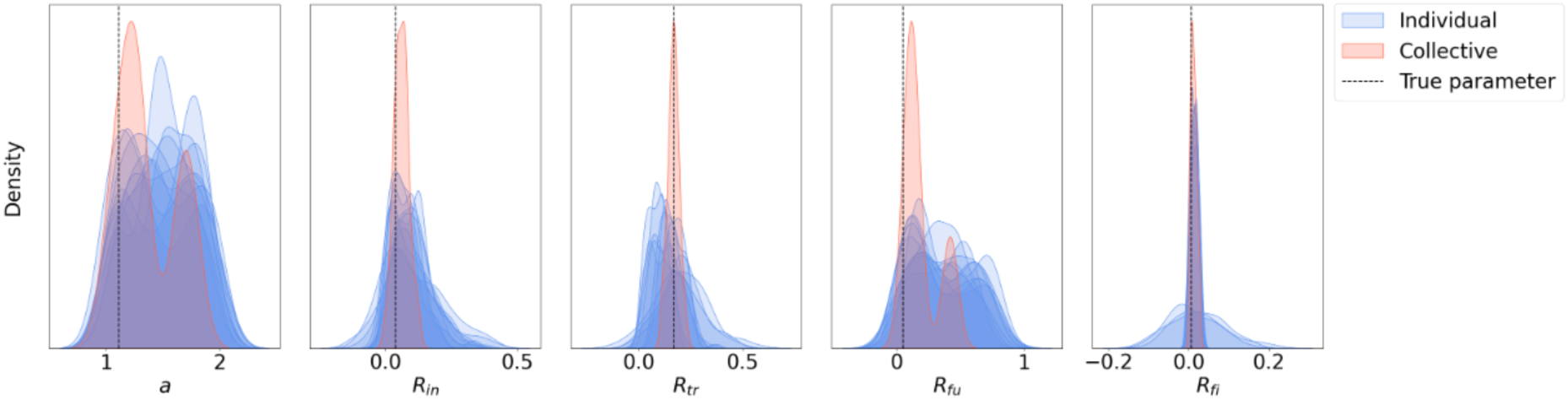
Collective posterior inference from genome rearrangement model simulations. Marginal individual posterior distributions of ten synthetic observations, inferred using rejection-ABC (blue); marginal collective posterior distributions, aggregated from the individual posterior distributions (red); and the true parameter values (black dotted line) for the five model parameters.

**Figure S12.**
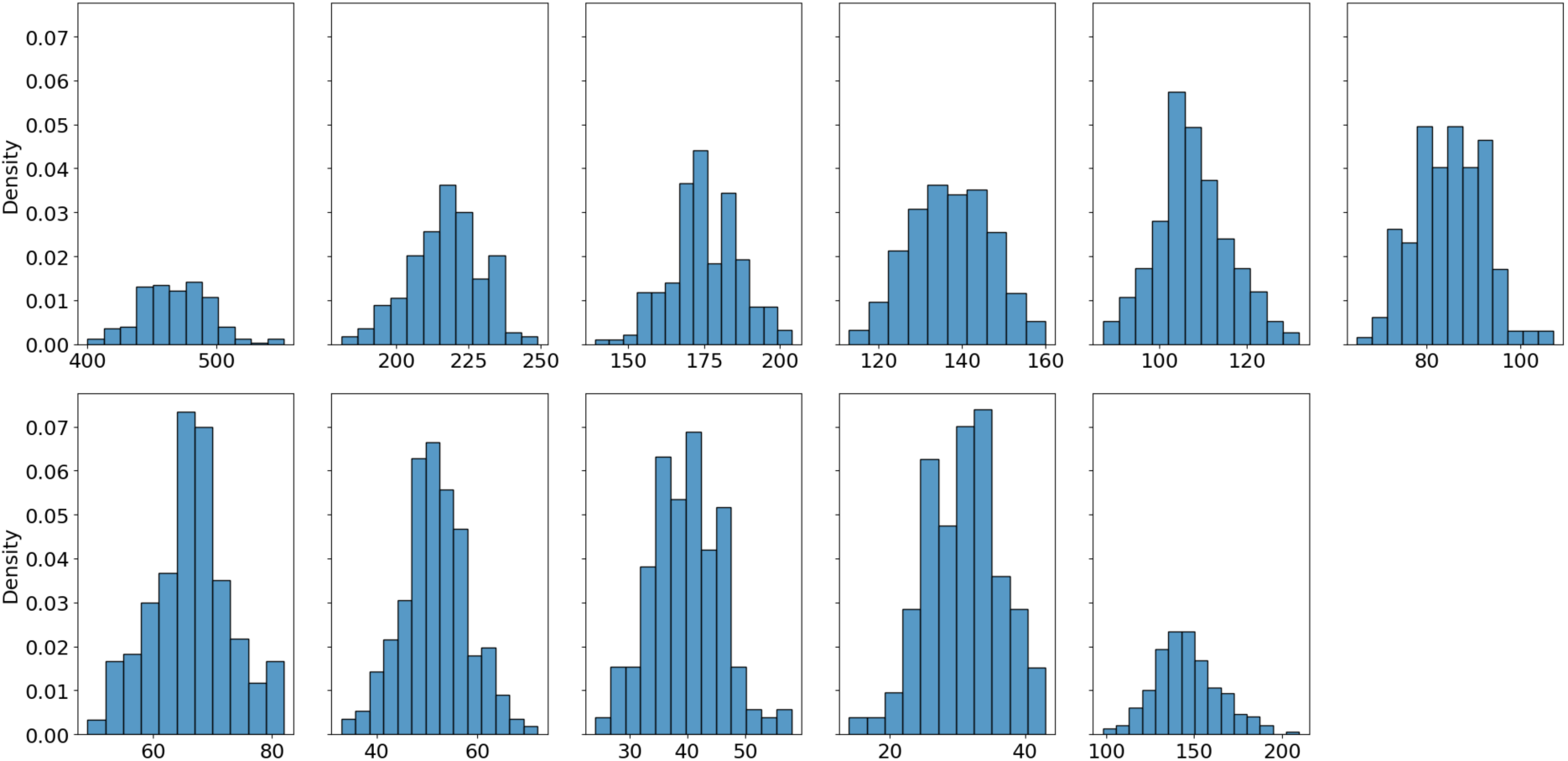
Distribution of phylogenetic summary statistics. Histograms of summary statistics from 100 simulations of the estimated parameter values from Moshe et al. (2022).

### Additional Results – Synthetic Evolutionary Benchmark

**Figure S13.**
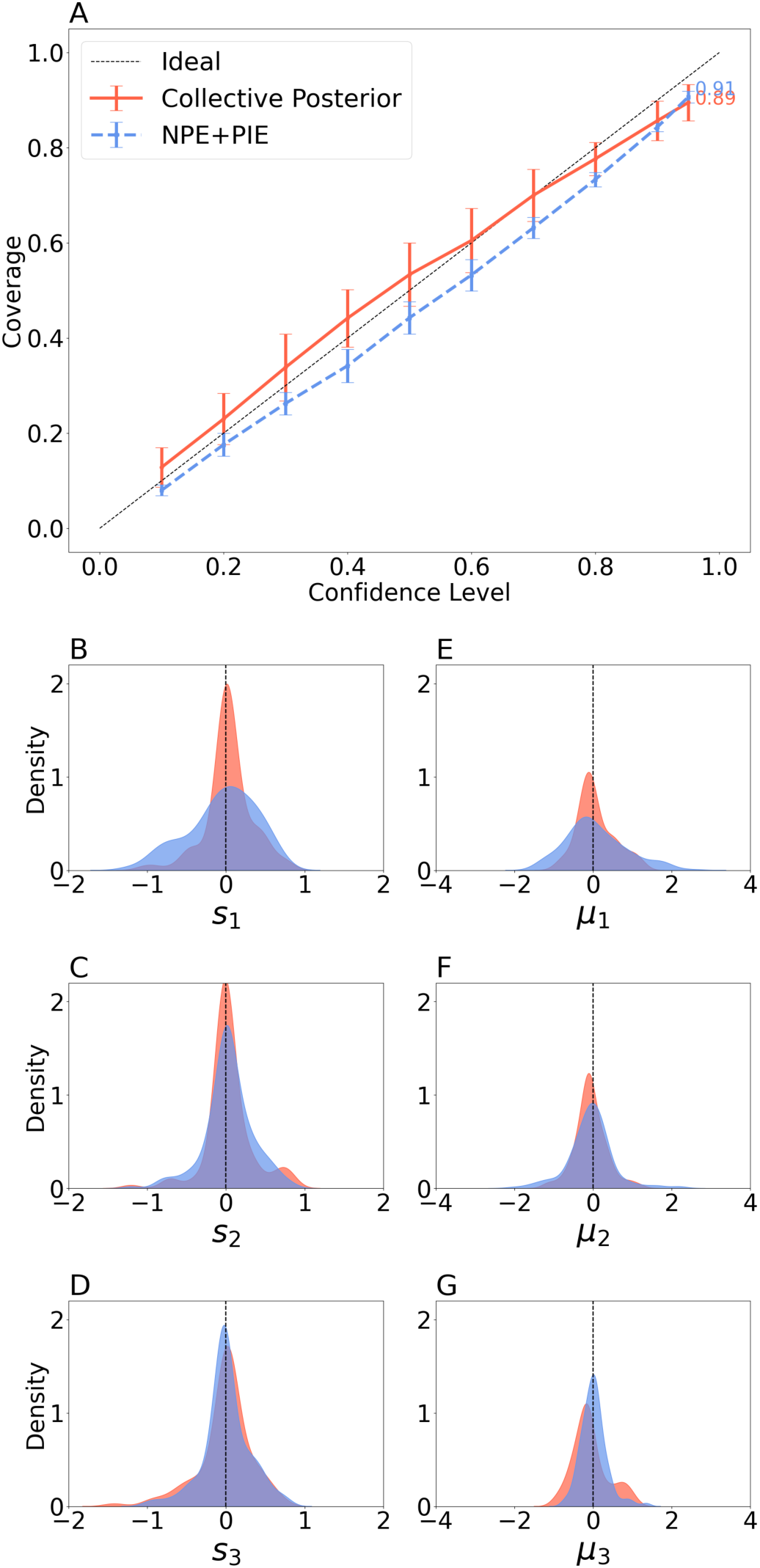
The collective posterior with NPE and NPE+PIE are comparable on clean evolutionary simulations. **(A) Coverage plots.** Box plots of coverages Pr[*θ*_*i*_ ∈ *HDR*_*α*_], corresponding to confidence levels *α*. **(B-G) Centrality.** Distributions of the difference between the posterior sample mean and the true parameter for all model parameters.

**Figure S14.**
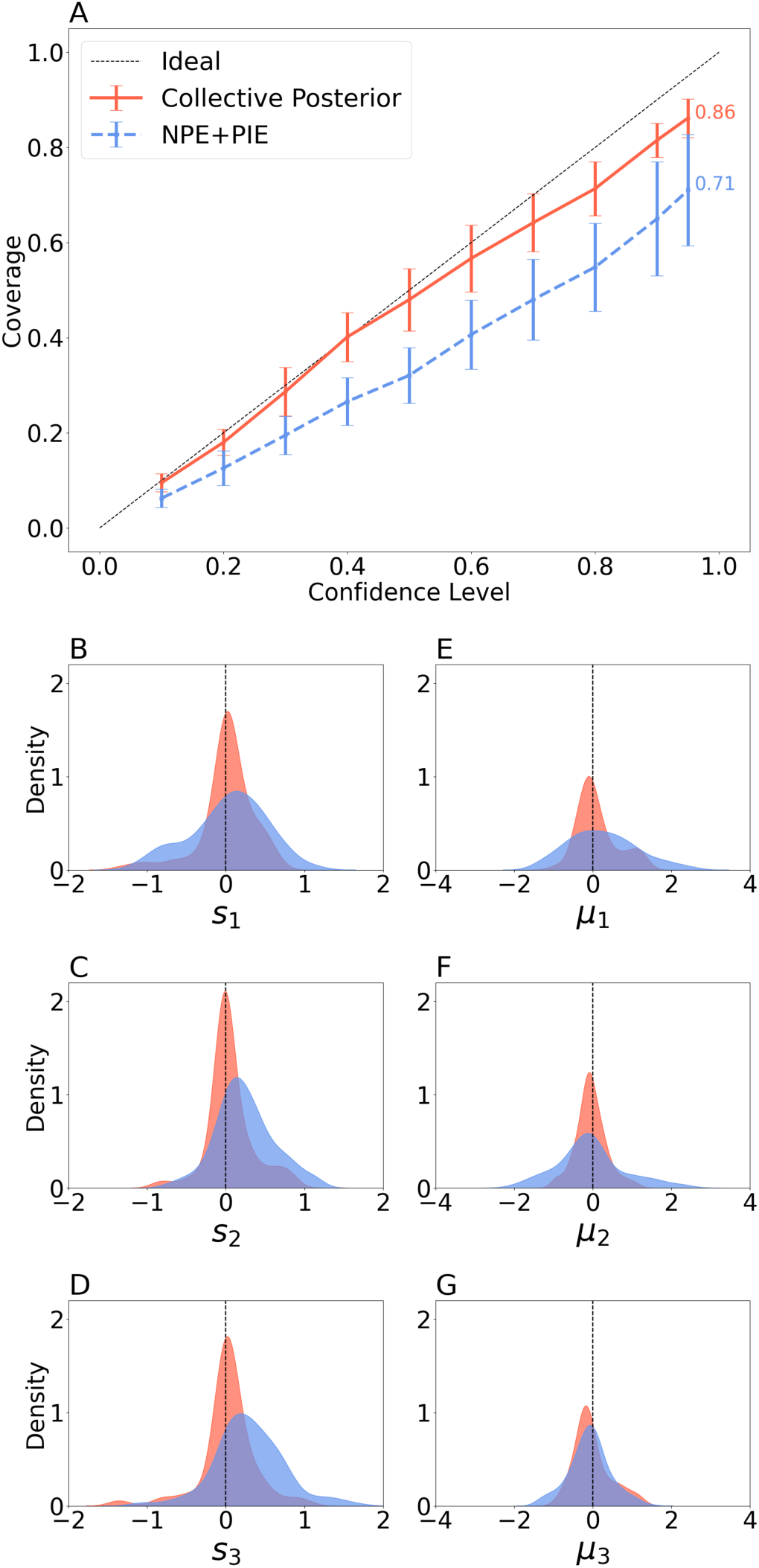
The collective posterior with NPE outperforms NPE+PIE on hierarchical evolutionary simulations (without measurement noise). **(A) Coverage plots.** Box plots of coverages Pr[*θ*_*i*_ ∈ *HDR*_*α*_], corresponding to confidence levels *α*. **(B-G) Centrality.** Distributions of the difference between the posterior sample mean and the true parameter for all model parameters.

### Single-locus Wright–Fisher task

We evaluated the collective posterior distribution and NPE+PIE on a Wright–Fisher simulator tracking a single locus with one beneficial mutation. The model follows the same evolutionary process as the three-locus setup (mutation → selection → drift), but observations consist of 20 samples recorded every 10 generations. In this setting, we infer the selection coefficient *s*, mutation rate *δ*, and effective population size *N*_*e*_. Although this simulator is computationally efficient, it lacks inherent variance that would necessitate collective inference methods (Figure S15). When tested on ten replicate observations, NPE+PIE showed a clear advantage in estimating the selection coefficient, whereas the collective posterior was more accurate for *N*_*e*_ (Figure S16). However, after perturbing the parameters with Gaussian noise *N*(0,0.2), the collective posterior outperformed NPE+PIE across all parameters (Figure S17).

**Figure S15.**
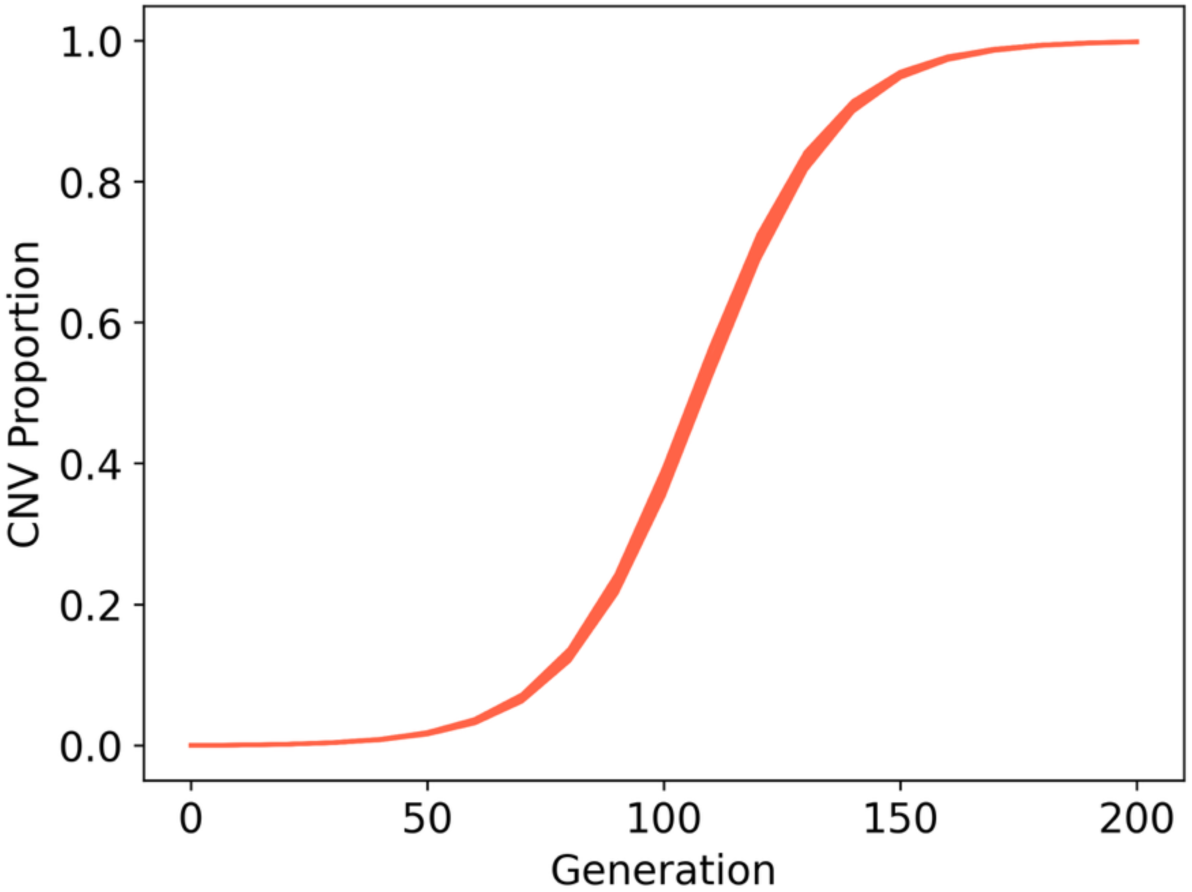
Simulations from the single-locus WF model have an extremely low variance. 10 replicates of the same parameter values fed to the single-locus WF simulator (random seed, unfixed).

**Figure S16.**
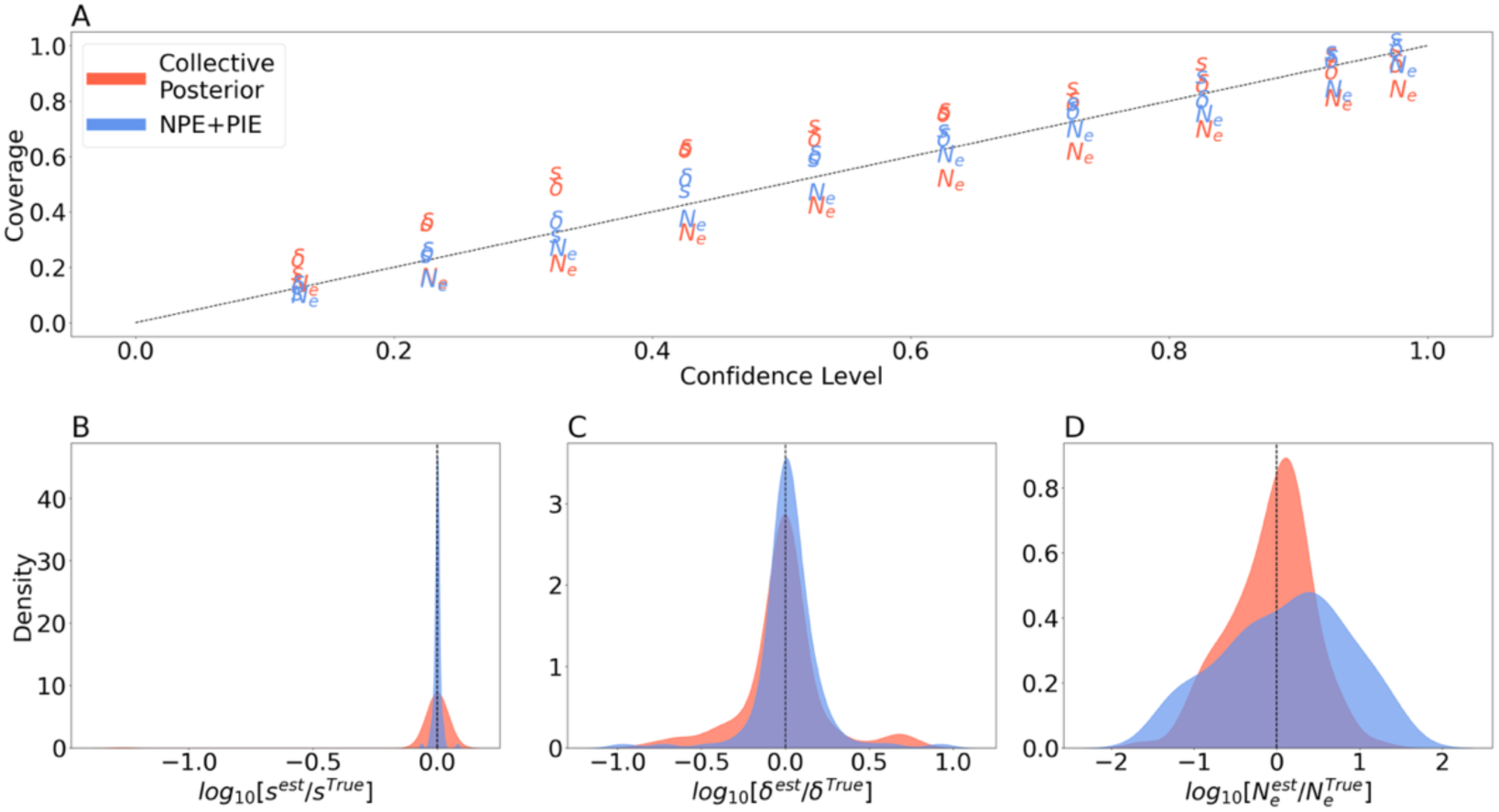
NPE+PIE and the collective posterior are comparable on clean single-locus WF simulations. **(A) Coverage plots.** Box plots of coverages Pr[*θ*_*i*_ ∈ *HDR*_*α*_], corresponding to confidence levels *α*. **(B-D) Centrality.** Distributions of the difference between the posterior sample mean and the true parameter for all model parameters. NPE+PIE shows dranatically better results for the selection coefficient *s*, however underpeforms for *N*_*e*_.

**Figure S17.**
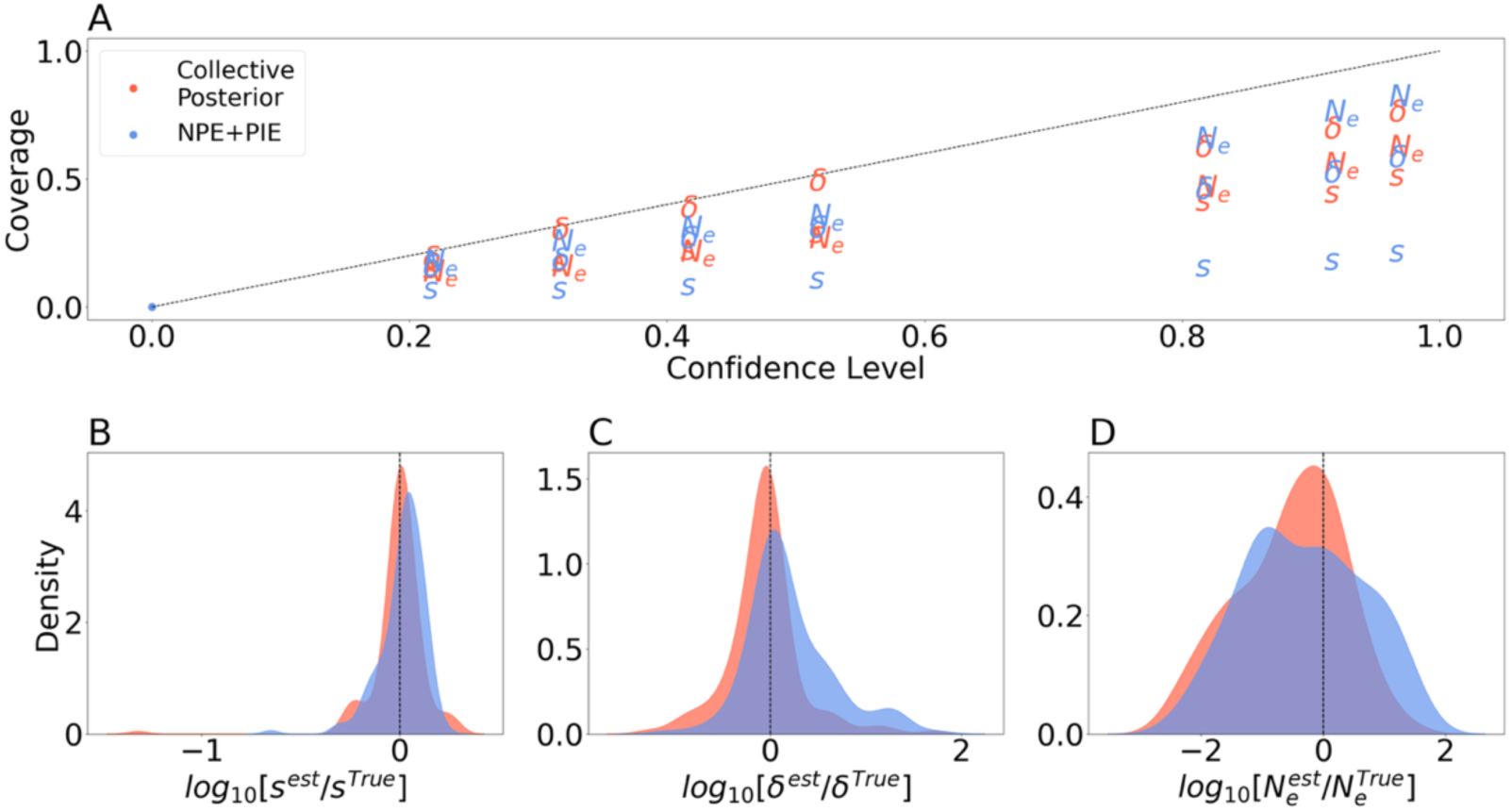
The collective posterior with NPE outperforms NPE+PIE on perturbed single-locus WF simulations. **(A) Coverage plots.** Box plots of coverages Pr[*θ*_*i*_ ∈ *HDR*_*α*_], corresponding to confidence levels *α*. **(B-D) Centrality.** Distributions of the difference between the posterior sample mean and the true parameter for all model parameters.

### Floor-Raising Ensembles

If we view the collective posterior distribution as a robust posterior aggregator, we can apply it to single observations in the context of ensembles. Much like experimental outliers, one or more of the models in the ensembles could be biased or overconfident. This will either skew or overspread the posterior distribution. In contrast, using the collective posterior could produce an ensemble that concentrates on the intersection of high-density regions (Figure S18).

**Figure S18.**
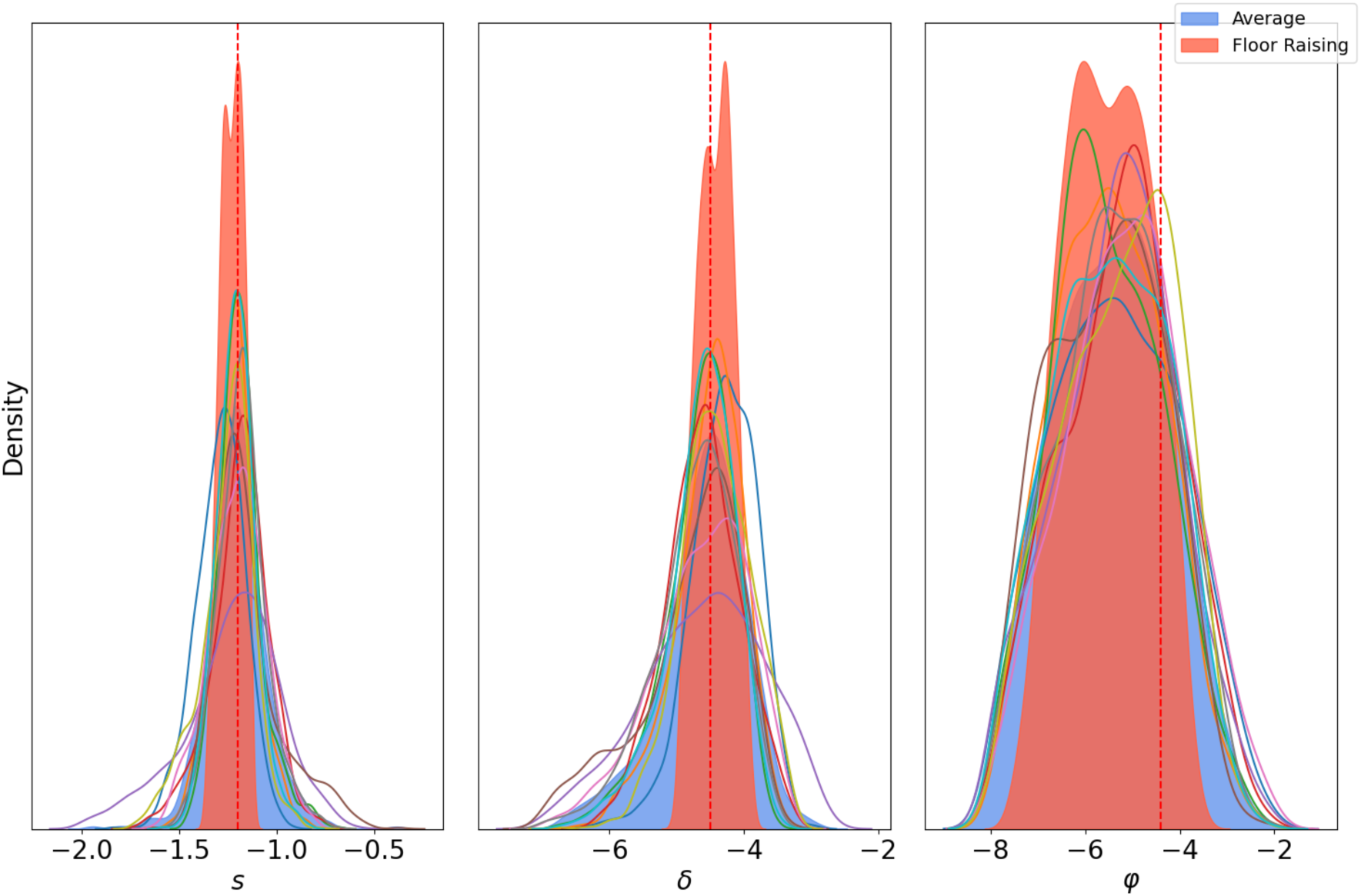
Improved precision via “Floor Raising” ensemble. To assess the impact of ensembling on estimator accuracy, we trained 10 independent NPE neural density estimators on the same Wright-Fisher task and evaluated them on a single replicate. Thin lines show the posterior distributions from each of the 10 density estimators, illustrating stochastic variation in neural estimator training. Blue distribution represents the standard averaging ensemble (arithmetic mean of the posterior), which broadens the posterior to encompass the variance of all individual estimators. Red distribution represents the proposed “Floor Raising” ensemble, calculated using the robust collective posterior formula (Eq. 7) across the 10 estimators. By treating each estimator as independent experts, the Floor Raising method identifies the consensus region where all estimators agree, resulting in significantly sharper parameter estimates compared to averaging ensemble.

### Benchmarking Wall Times

**Table S1.** The Collective Posterior is faster compared to NPE+PIE and NPSE. Training (including simulation) and inference times for all methods, tested on the Chuong et al. (2025) task. While NPE+PIE offers rapid sampling, the Collective Posterior requires significantly less training time, with a small inference overhead. In addition, NPSE needs a short training time, but a significantly longer inference time. Analysis performed on 48 CPU cores with NPE and NPSE (5) default hyperparameters. Inference corresponds to 10,000 samples from the estimated posterior for each of the 500 observation sets. For the collective posterior, we report the running times for both sampling strategies: SIR (sampling importance resampling), and MCMC (Markov Chain Monte Carlo).

| Method | Simulations | Training + simulation<br>(hours) | Inference<br>(seconds) | Total time<br>(hours) |
| --- | --- | --- | --- | --- |
| Collective<br>Posterior (NPE) | 10K | 0.1 | 0.53 (SIR)<br>102 (MCMC) | 0.17 (SIR)<br>14.26 (MCMC) |
| NPE + PIE | 10K | 0.99 | 0.12 | 1.01 |
| Collective<br>Posterior (NPE) | 30K | 0.33 | 0.53 (SIR)<br>102 (MCMC) | 0.4 (SIR)<br>14.49 (MCMC) |
| NPE + PIE | 30K | 3.5 | 0.12 | 3.52 |
| NPSE | 30K | 0.34 | 117 | 16.59 |

**Table S2.**
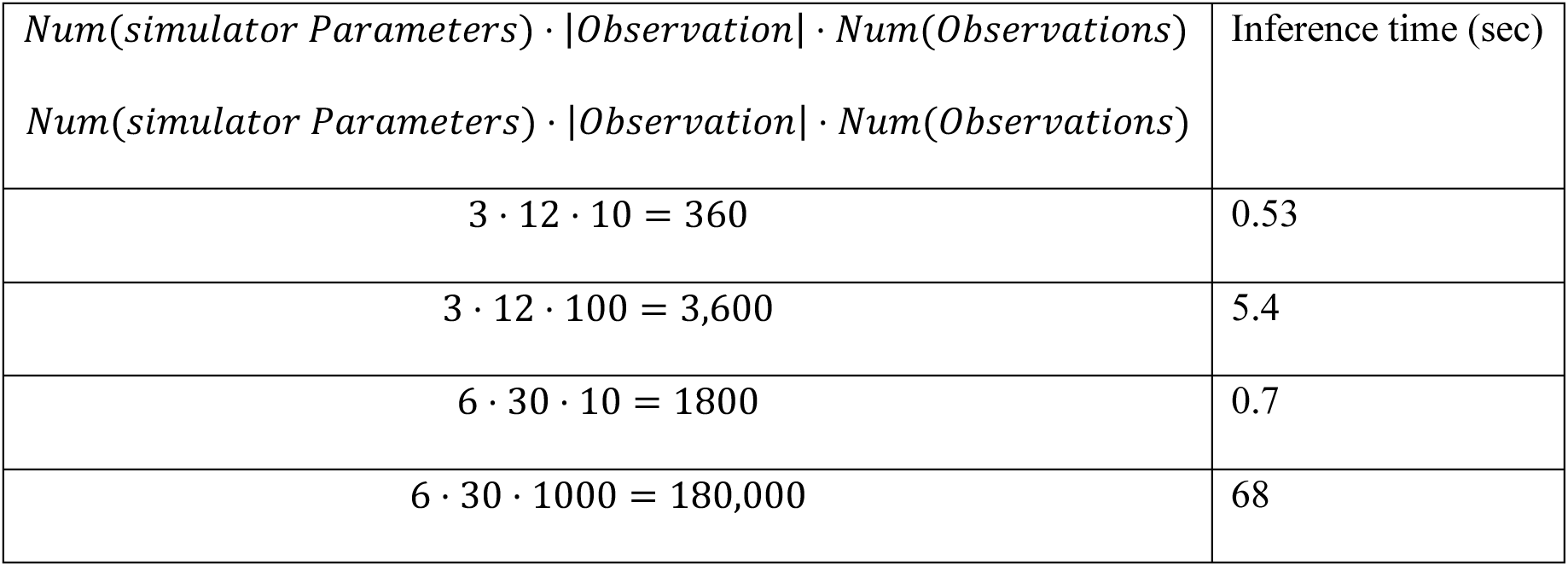
Scalability of the collective posterior with importance sampling. The collective posterior can easily scale to 100 or even 1,000 observations in reasonable times, while using the same neural density estimator that required less than 1 hour to train. For comparison, training NPE+PIE for datsets of 1,000 observations would take about two weeks, and inference of NPSE would take about 36 hours.

|  |  |
| --- | --- |
| $Num(simulator\ Parameters) \cdot \vert Observation\vert \cdot Num(Observations)$ | Inference time (sec) |
| $Num(simulator\ Parameters) \cdot \vert Observation\vert \cdot Num(Observations)$ | |
| $3 \cdot 12 \cdot 10 = 360$ | 0.53 |
| $3 \cdot 12 \cdot 100 = 3,600$ | 5.4 |
| $6 \cdot 30 \cdot 10 = 1800$ | 0.7 |
| $6 \cdot 30 \cdot 1000 = 180,000$ | 68 |

### Erratic Posterior Densities

**Table S3.** Erratic posterior densities. Log-posterior probability by NPE of both prior edges and the sample mean, conditioned on the empirical data shown in supplementary Figure S4. While the posterior density is consistent for a given observation, the densities for unlikely values change by hundreds of orders of magnitude among observations. The density of the sample mean is similar among observations.

| Observation ($i$) | $\log [p(\theta_{min}\vert x_i)]$ | $\log [p(\theta_{max}\vert x_i)]$ | $\log [p(\bar{\theta}\vert x_i)]$ |
| --- | --- | --- | --- |
| 1 | -1394 | -1394 | 4 |
| 2 | -2369 | -2369 | 4 |
| 3 | -1407 | -1407 | 3 |
| 4 | -3653 | -3653 | 5 |
| 5 | -2273 | -2273 | 4 |
| 6 | -2656 | -2656 | 4 |
| 7 | -3305 | -3305 | 5 |

